# MATCAP1 preferentially binds an expanded tubulin conformation to generate detyrosinated and ΔC2 α-tubulin

**DOI:** 10.1101/2025.08.14.670257

**Authors:** Yang Yue, Takashi Hotta, Ryoma Ohi, Kristen J. Verhey

## Abstract

Microtubules are cytoskeletal filaments with critical roles in cell division, cell motility, intracellular trafficking, and cilium function. In cells, subsets of microtubules are selectively marked by posttranslational modifications (PTMs), which control the ability of microtubule-associated proteins (MAPs) and molecular motors to engage microtubules. Detyrosination (ΔY) and ΔC2 are PTMs of α-tubulin, wherein one or two residues, respectively, are enzymatically removed from the C-terminus of the protein. How specific patterns of PTMs are generated in cells is not understood. Here, we use in vitro reconstitution assays to investigate the microtubule binding behavior of metallopeptidase MATCAP1 and the mechanism by which it generates ΔY and ΔC2 modifications of α-tubulin. We demonstrate that MATCAP1 preferentially binds to microtubules composed of tubulin subunits in an expanded conformation, which can be induced by preventing β-tubulin GTP hydrolysis, Taxol treatment, or kinesin-1 stepping. MATCAP1 binds to expanded microtubule lattices with long dwell time and sequentially removes the terminal tyrosine residue to generate ΔY-microtubules and the penultimate glutamate residue to generate ΔC2-microtubules. Thus, the lattice conformation of microtubules is a key factor that gates the binding and activity of MATCAP1.

## Introduction

Microtubules are dynamic polymers of α,β-tubulin that play critical roles in eukaryotic cells, ranging from cell division to cell migration and intracellular trafficking. α,β-tubulin is a GTPase, and the nucleotide-bound state of β-tubulin is tied to the ability of microtubules to cycle between states of assembly and disassembly (Cleary and Hancock 2021, Gudimchuk and McIntosh 2021, Chew and Cross 2025). In the GTP-bound form, tubulin readily assembles into microtubules. Upon incorporation into the microtubule lattice, β-tubulin hydrolyzes its bound nucleotide, generating GDP-bound β-tubulin. Cryo-electron microscopy studies show that GTP-tubulin is lengthened (“expanded”) by ∼1-2 Å along the longitudinal axis of the microtubule lattice relative to GDP-tubulin (“compacted”) (Alushin et al. 2014, Manka and Moores 2018, Zhang et al. 2018, LaFrance et al. 2022). These nucleotide-dependent conformational states are thought to impact tubulin-tubulin interactions in the microtubule lattice and thereby affect microtubule dynamics.

In cells, the functional properties of microtubules are altered through post-translational modifications (PTMs) of their tubulin subunits, and these in turn act by changing the cohort of microtubule-associated proteins (MAPs) that interact with microtubules (Verhey and Gaertig 2007, Janke and Magiera 2020, Roll-Mecak 2020). An important class of tubulin PTMs consists of those that involve proteolytic removal of amino acids from the α-tubulin C-terminal tail (αCTT): detyrosinated (ΔY)-, ΔC2-, and ΔC3-α-tubulin. Of these, microtubule detyrosination is the best understood in terms of its physiological roles and effects on MAPs (Nieuwenhuis and Brummelkamp 2019, Ferreira et al. 2020, Moutin et al. 2021, Sanyal et al. 2023). For example, ΔY-microtubules near the poles of spindle microtubules facilitates chromosome congression by the kinesin-7 CENP-E (KIF10) whereas unmodified microtubules (tyrosinated or Y-microtubules) near the kinetochores enable proper kinesin-13 MCAK (KIF2C) error correction (Peris et al. 2009, Barisic et al. 2015, Liao et al. 2019). In interphase cells, the Y/ΔY state regulates intracellular trafficking as ΔY-microtubules are preferentially utilized by kinesin-1 whereas Y-microtubules are preferentially utilized by cytoplasmic dynein-1 (Dunn et al. 2008, Cai et al. 2009, Konishi and Setou 2009, McKenney et al. 2016, Nirschl et al. 2016, Lavrsen et al. 2023, Konietzny et al. 2024). The Y/ΔY state also affects trafficking in cilia where ΔY-microtubules promote the anterograde transport of intraflagellar transport trains by kinesin-2 and Y-microtubules enable the dynein-2-dependent retrograde trafficking of trains (Chhatre et al. 2025). In cardiomyocytes, microtubule detyrosination promotes a physical linkage between microtubules and sarcomeres, creating a force-restoring system that opposes myocyte contraction (Robison et al. 2016, Chen et al. 2018, Salomon et al. 2022).

How cells select microtubules for detyrosination or decide where to generate ΔC2- and ΔC3-α-tubulin has been a mystery for decades. Recent work has revealed that vertebrates encode two families of microtubule detyrosination enzymes: 1) the vasohibins (VASH1 and VASH2), cysteine proteases which require <u>s</u>mall-<u>v</u>asohibin <u>b</u>inding <u>p</u>rotein (SVBP) as a cofactor (Aillaud et al. 2017, Nieuwenhuis et al. 2017) and 2) the microtubule-associated tyrosine carboxypeptidases (MATCAP1 and MATCAP2), metalloproteases also known as tubulin metallocarboxypeptidases (TMCP1 and TMCP2) (Landskron et al. 2022, Nicot et al. 2023) [reviewed in (Bak et al. 2024)]. Deletion of VASH1/2 and MATCAP1 proteins in mice and various cell lines abolishes the production of ΔY-microtubules (Landskron et al. 2022), suggesting that they are the only enzymes capable of microtubule detyrosination activity in mammals.

The writers of ΔC2- and ΔC3-a-tubulin marks have long been assumed to be cytosolic carboxypeptidases (CCPs) as these enzymes can generate ΔC2- and ΔC3-a-tubulin when overexpressed in mammalian cells (Rogowski et al. 2010, Berezniuk et al. 2012, Tort et al. 2014, Aillaud et al. 2016). Consistent with this possibility, we previously showed that CCP1 is the primary enzyme that generates ΔC2-a-tubulin in HeLa cells lacking tubulin tyrosine ligase (TTL), an enzyme that converts ΔY-a-tubulin back to full-length (tyrosinated) a-tubulin (Hotta et al. 2023). However, tissues from *pcd* mice, which harbor an inactivating mutation in CCP1, retain high levels of ΔC2-a-tubulin, suggesting that additional enzyme(s) can generate ΔC2-a-tubulin (Rogowski et al. 2010). Interestingly, recent work demonstrated that MATCAP1/TMCP1 and TMCP2 are both capable of generating ΔC2-a-tubulin in addition to ΔY-α-tubulin (Nicot et al. 2023).

In previous work, we developed a microscopy-based enzymatic assay to track the generation of ΔY-microtubules by VASH1/SVBP in real time. We discovered that VASH1/SVBP binds to microtubules regardless of their nucleotide or conformational state. However, the ability of VASH1/SVBP to detyrosinate microtubules is dramatically enhanced when tubulin subunits are in an expanded conformation, *i.e.*, on microtubules assembled with the non-hydrolyzable analogue GMPCPP or GDP-microtubules that are stabilized with the drug Taxol (Yue et al. 2023). These findings provide an explanation for why most microtubules in cells, which exist in a GDP-bound or compacted state, are not selected for modification by the VASH1/SVBP enzyme.

In this work, we sought to extend our understanding of the relationship between the conformational state of the microtubule lattice and detyrosination by examining the properties of MATCAP1. We find that, unlike VASH1/SVBP, MATCAP1 preferentially binds to microtubules with tubulin subunits in an expanded conformation, which can be induced by preventing β-tubulin GTP hydrolysis, Taxol treatment, or stepping of kinesin-1 along the microtubule lattice. Once bound, MATCAP1 exhibits a long dwell time, in contrast to VASH1/SVBP which undergoes brief interactions with the microtubule lattice. Using probes that specifically recognize ΔY- and ΔC2-microtubules and the microscopy-based enzymatic assay, we show that MATCAP1’s rate of generating ΔY-microtubules is faster than for generating ΔC2-microtubules. This work provides a molecular explanation of how MATCAP1 sequentially generates ΔY- and ΔC2-a-tubulin within an expanded microtubule lattice.

## Results

### MATCAP1 directly generates both ΔY- and ΔC2-microtubules in HeLa cells

To investigate MATCAP1 activity in cells, we transiently expressed MATCAP1 tagged with both a PA epitope and TagRFP at the N-terminus (PA-TagRFP-MATCAP1) in HeLa Kyoto cells. No ΔY-microtubules were detected in the untransfected cells by immunostaining (**Figure 1A, asterisks**) or in cells expressing the control PA-TagRFP construct by western blot (**Figure 1C, Figure S1A**). However, transient expression of PA-TagRFP-MATCAP1 resulted in a dramatic increase in ΔY-microtubules by both immunostaining and western blot (**Figure 1, A and C; Figure S1A**). We also generated a HeLa Kyoto cell line that contains a stably-integrated cassette for expressing Halo-MATCAP1 in a doxycycline-inducible manner (Khandelia et al. 2011). In the absence of doxycycline, no ΔY-microtubules were detected, however, doxycycline-induced expression of Halo-MATCAP1 resulted in a concomitant increase in ΔY-microtubules (**Figure S1B**). These results are consistent with previous work showing that the enzymatic activity of MATCAP1 results in tubulin detyrosination (Landskron et al. 2022, Nicot et al. 2023).

**Figure 1.**
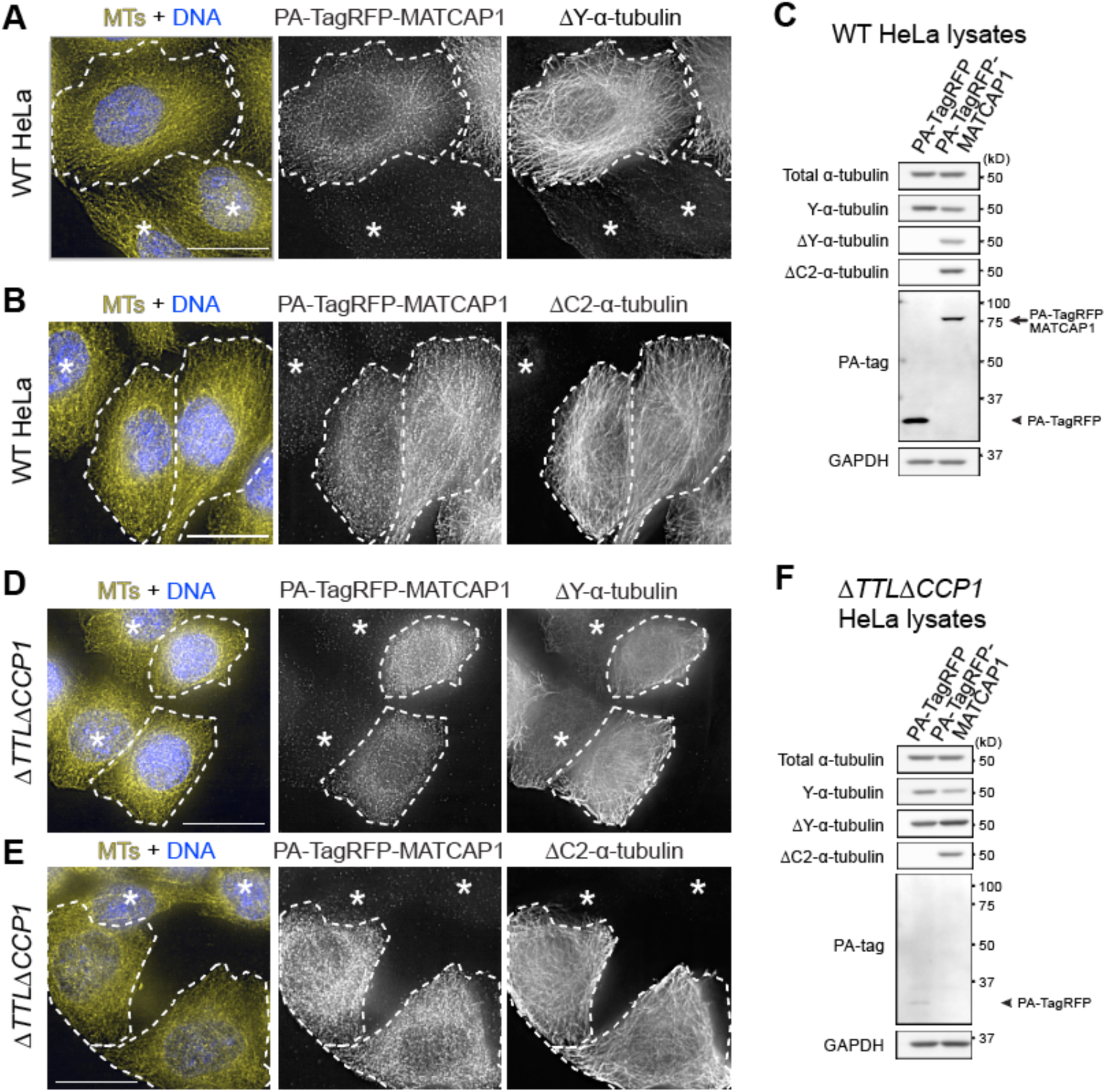
MATCAP1 directly generates both ΔY- and ΔC2-microtubules in cells. **(A-C)** Transient expression of MATCAP1 generates ΔY- and ΔC2-microtubules (MTs). **(A,B)** Representative images of HeLa cells transiently expressing PA-TagRFP-MATCAP1 and fixed and stained with antibodies against (**A**) ΔY-MTs or (**B**) ΔC2-MTs together with anti-PA-tag (to detect PA-TagRFP-MATCAP1) and AlexaFluor647-conjugated DM1a (to detect total MTs) antibodies. **(C)** Lysates from HeLa cells transiently expressing PA-TagRFP or PA-TagRFP-MATCAP1 were immunoblotted with the indicated antibodies. **(D-F)** MATCAP1 directly generates ΔC2-MTs in cells. **(D,E)** Representative images of Δ*TTL*Δ*CCP1* HeLa cells transiently expressing PA-TagRFP-MATCAP1 and fixed and stained with antibodies against (**D**) ΔY-MTs or (**E**) ΔC2-MTs together with anti-PA-tag (to detect PA-TagRFP-MATCAP1) and AlexaFluor647-conjugated DM1a (to detect total MTs) antibodies. **(F)** Lysates from Δ*TTL*Δ*CCP1* HeLa cells transiently expressing PA-TagRFP or PA-TagRFP-MATCAP1 were immunoblotted with the indicated antibodies. For all images, dashed lines indicate transfected cells, while asterisks indicate untransfected cells. Scale bars, 20 µm.

We next tested whether MATCAP1 activity results in increased levels of ΔC2-microtubules in cells. Indeed, we found that transient expression of PA-TagRFP-MATCAP1 (**Figure 1, B and C**) and doxycycline-induced expression of Halo-MATCAP1 (**Figure S1C**) in HeLa cells resulted in increased levels of ΔC2-microtubules, suggesting that MATCAP1 can carry out further trimming of the αCTT, in agreement with recent work (Nicot et al. 2023). ΔC2-microtubules in mammalian cells can also be generated by cytosolic carboxypeptidases (CCPs). While overexpression of five CCPs (CCP1, CCP2, CCP3, CCP4, and CCP6) can generate ΔC2-α-tubulin in cells, only loss of CCP1 decreases ΔC2-α-tubulin in cells and animals (Rogowski et al. 2010, Berezniuk et al. 2012, Tort et al. 2014, Hotta et al. 2023). To rule out the possibility that expression of MATCAP1 results in increased ΔC2-microtubules due to the activation of endogenous CCP1, we utilized a HeLa cell line (Δ*TTL*Δ*CCP1*) that lacks tubulin tyrosine ligase (TTL) and CCP1 proteins (Hotta et al. 2023). In the Δ*TTL*Δ*CCP1* cell line, knockout of TTL expression results in high levels of ΔY-microtubules (**Figure1D, asterisks; Figure 1F; Figure S1A**), while knockout of CCP1 prevents the formation of ΔC2-α-tubulin as assessed by immunofluorescence (**Figure 1E, asterisks**) and western blot (**Figure 1F; Figure S1A**) (Hotta et al. 2023). Overexpression of MATCAP1 in Δ*TTL*Δ*CCP1* cells produced detectable ΔC2-microtubules, as assessed by both assays (**Figure 1, E and F; Figure S1, A and C**), indicating that MATCAP1 can directly generate both ΔY-microtubules and ΔC2-microtubules, consistent with recent work (Nicot et al. 2023).

### MATCAP1 directly generates both ΔY-microtubules and ΔC2-microtubules in vitro

To investigate the catalytic properties of MATCAP1, we utilized a total internal reflection fluorescence (TIRF) microscopy-based enzymatic assay developed in our previous work on VASH1/SVBP (Yue et al. 2023). We purified tubulin from HeLa S3 cells as they lack most PTMs, including ΔY-tubulin (Souphron et al. 2019, Thomas et al. 2025), and expressed full-length human MATCAP1 tagged with Halo tag at the N-terminus (Halo-MATCAP1) or VASH1-Halo with SVBP in COS-7 cells. Taxol-stabilized HeLa microtubules were adhered to the surface of flow chambers and then incubated with cell lysates containing either MATCAP1 or VASH1/SVBP (unlabeled) enzymes together with a fluorescently-labeled probe for detecting the modified tubulin (**Figure 2A**). We first examined the generation of ΔY-microtubules. As a probe, we used a fluorescently-labeled (Alexa488) fragment antibody binding (Fab) generated from the RM444 recombinant monoclonal antibody that specifically recognizes ΔY-α-tubulin [ΔY-Fab^488^, (Yue et al. 2023)]. Incubation of HeLa microtubules with either MATCAP1 or VASH1/SVBP for 10 min resulted in a dramatic increase in the amount of ΔY-Fab^488^ probe bound to microtubules (**Figure 2B**). To further examine the activities of MATCAP1 and VASH1/SVBP, we added 1 nM of each protein in cell lysate and observed their enzymatic activities by time-lapse imaging. We found that both enzymes display strong detyrosination activity (**Figure 2, C and D**).

**Figure 2.**
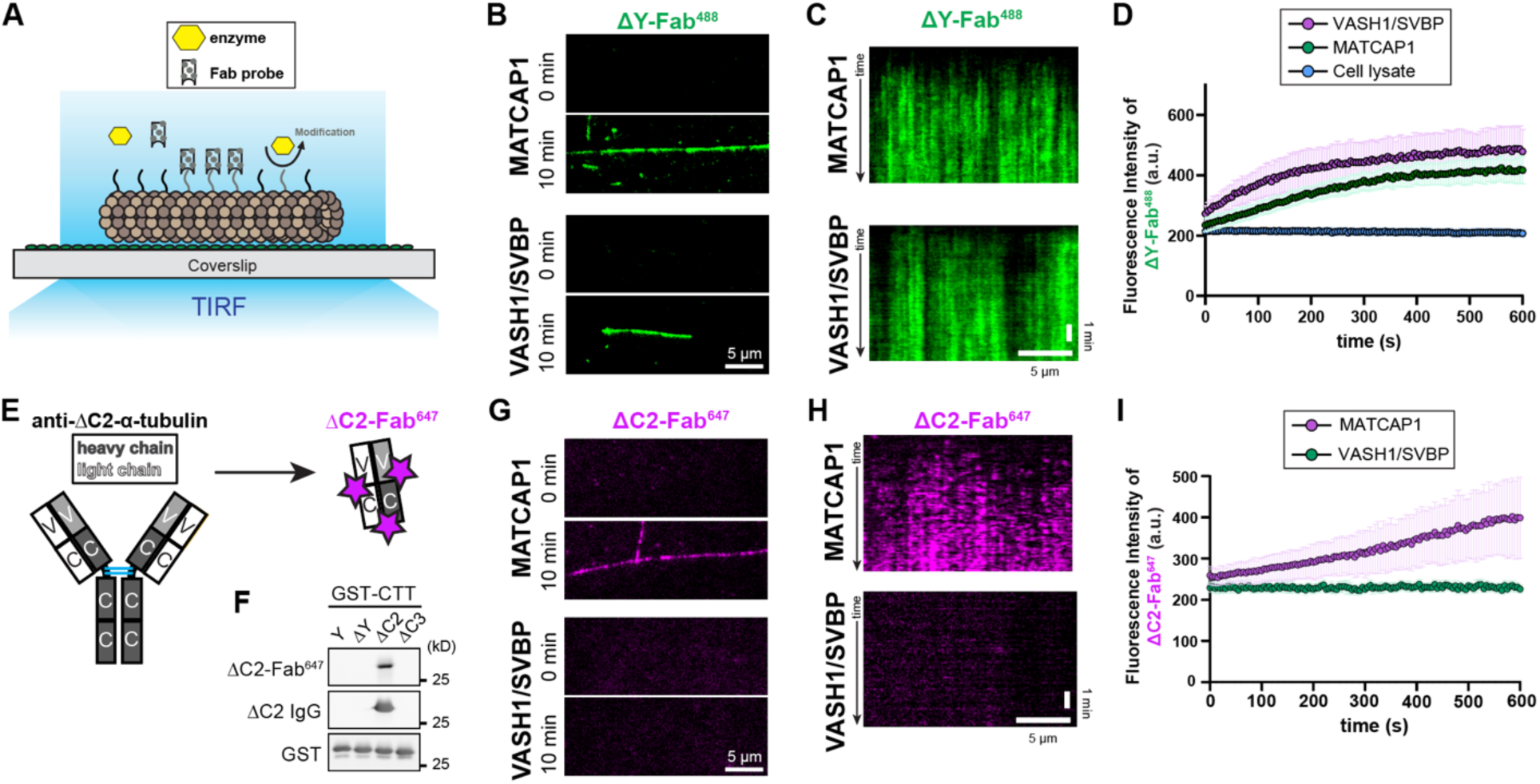
MATCAP1 directly generates both ΔY- and ΔC2-MTs in vitro. **(A)** Schematic of the microscopy-based enzymatic assay for detecting the biogenesis of microtubule PTMs. Taxol-stabilized HeLa MTs were incubated with a modifying enzyme (yellow) and a fragment antibody binding (Fab) probe (grey) that recognizes a specific tubulin PTM. Binding of the Fab probe to microtubules is observed by total internal reflection fluorescence (TIRF) microscopy. **(B-D)** Biogenesis of ΔY-MTs. (**B**) Representative images of ΔY-Fab^488^ labeling of microtubules at 0 min and after 10 min incubation with 1 nM Halo-MATCAP1 or VASH1/SVBP-Halo in cell lysates. Scale bar, 5 µm. **(C)** Representative kymographs showing ΔY-Fab^488^ probe labeling of microtubules over time after the addition of 1 nM Halo-MATCAP1 or VASH1/SVBP-Halo in cell lysates. Time is shown on the y-axis (scale bar, 1 min), and distance along the microtubule is on the x-axis (scale bar, 5 µm). **(D)** Quantification of the mean fluorescence intensity of ΔY-Fab^488^ probe labeling along microtubules over time for the experiments in (C). Data are presented as mean ± S.D., with n = 26–41 microtubules from two independent experiments. **(E,F)** Development of a Fab probe for ΔC2-MTs. **(E)** Schematic showing the RM447 monoclonal antibody against ΔC2-α-tubulin. The antibody was non-specifically labeled with Alexa 647 dye and then a Fab fragment (ΔC2-Fab^647^) was produced by papain digestion. **(F)** Purified glutathione S-transferase (GST) proteins fused to various α-tubulin C-terminal tail (CTT) sequences were analyzed by western blot using the ΔC2-Fab^647^ probe (top), the RM447 monoclonal ΔC2-α-tubulin antibody (middle), or an antibody against GST (bottom). The α-tubulin CTT sequences are full-length CTT (Y), missing the C-terminal tyrosine (ΔY), missing the C-terminal two amino acid residues (ΔC2), and missing the C-terminal three amino acid residues (ΔC3). **(G-I)** Biogenesis of ΔC2-MTs. (**G**) Representative images of ΔC2-Fab^647^ labeling of microtubules at 0 min and after 10 min incubation with 1 nM Halo-MATCAP1 or VASH1/SVBP-Halo in cell lysates. Scale bar, 5 µm. **(H)** Representative kymographs showing ΔC2-Fab^647^ probe labeling of microtubules over time after the addition of 1 nM Halo-MATCAP1 or VASH1/SVBP-Halo in cell lysates. Time is shown on the y-axis (scale bar, 1 min), and distance along the microtubule is on the x-axis (scale bar, 5 μm). **(I)** Quantification of the mean fluorescence intensity of ΔC2-Fab^647^ probe labeling along microtubules over time for the experiments shown in (H). Data are presented as mean ± S.D., with n = 17–39 microtubules from two to four independent experiments.

To examine the generation of ΔC2-microtubules in real time, we generated a Fab version of the RM447 recombinant monoclonal antibody that specifically recognizes ΔC2-microtubules (Hotta et al. 2023). The ΔC2-Fab was fluorescently labeled with Alexa647 dye to generate a fluorescent ΔC2-Fab^647^ probe (**Figure 2E**). We confirmed that the ΔC2-Fab^647^ probe shows the same selectivity for ΔC2-microtubules as the original monoclonal antibody by western blotting (**Figure 2F**). Incubation of Taxol-stabilized HeLa microtubules and the ΔC2-Fab^647^ probe with cell lysate containing MATCAP1 for 10 min resulted in a dramatic increase in ΔC2-Fab^647^ probe binding to microtubules (**Figure 2G**). In contrast, incubation of HeLa microtubules and the ΔC2-Fab^647^ probe with cell lysate containing VASH1/SVBP for 10 min did not increase ΔC2-Fab^647^ probe binding to microtubules (**Figure 2G**). Using time-lapse imaging of assays containing 1 nM of MATCAP1 or VASH1/SVBP proteins in cell lysates, we found that only MATCAP1 activity resulted in an increase in ΔC2-microtubules over time (**Figure 2, H and I**). Taken together, these results demonstrate that only MATCAP1 is capable of generating both ΔY-microtubules and ΔC2-microtubules *in vitro*, consistent with our observations in cells.

### MATCAP1 generates ΔY-microtubules faster than ΔC2-microtubules

We next used the ΔY-Fab^488^ and ΔC2-Fab^647^ probes to directly compare the ability of MATCAP1 to generate the ΔY and ΔC2 modifications. Taxol-stabilized HeLa microtubules were incubated with cell lysate containing 1 nM MATCAP1 protein in the presence of both the ΔY-Fab^488^ and the ΔC2-Fab^647^ probes (**Figure 3A**). After a 10 min incubation, both ΔY-Fab^488^ and ΔC2-Fab^647^ probes were observed to decorate the same microtubules (**Figure 3B**). We thus carried out time-lapse imaging and found that the ΔY-Fab^488^ probe rapidly labeled the microtubule whereas the labeling by the ΔC2-Fab^647^ probe appeared more slowly and was more sparse (**Figure 3C**). Quantification of these assays showed that the ΔY-Fab^488^ fluorescence intensity plateaued 6–8 min after adding 1 nM MATCAP1, whereas the ΔC2-Fab^647^ intensity continued to increase throughout the 10 min imaging period (**Figure 3D).** With extended imaging time and a higher concentration of MATCAP1, we observed that the binding of the ΔC2-Fab^647^ probe to microtubules steadily increased over 1 hr of imaging (**Figure S2**). These results suggest that MATCAP1 generates ΔY-microtubules faster than ΔC2-microtubules.

**Figure 3.**
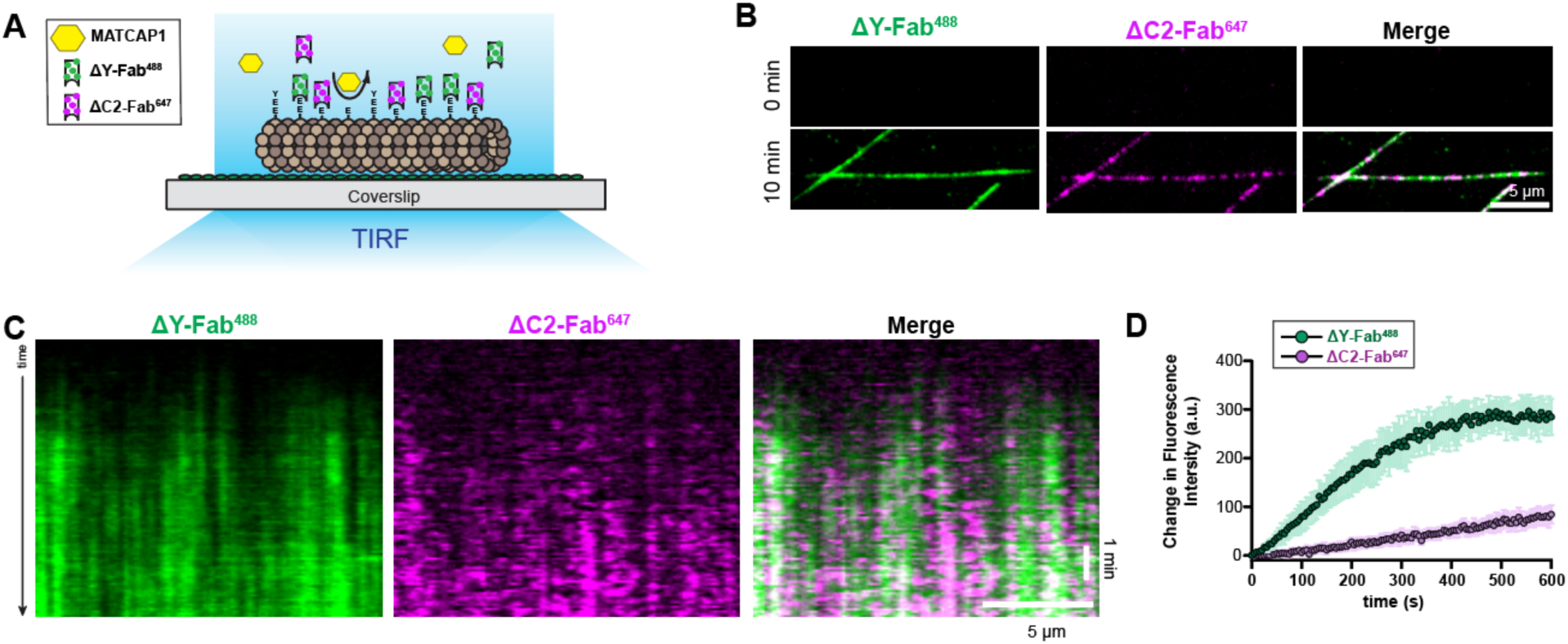
MATCAP1 generates ΔY-MTs faster than ΔC2-MTs in vitro. **(A)** Schematic of the microscopy-based enzymatic assay for dual Fab probe visualization of PTM biogenesis. Taxol-stabilized HeLa MTs were incubated with MATCAP1 and the formation of ΔY-MTs and ΔC2-MTs was observed by Fab binding to the microtubules. (**B**) Representative images of ΔY-Fab^488^ (green) and ΔC2-Fab^647^ probe (magenta) labeling of the same microtubules at 0 min and after 10 min incubation with 1 nM Halo-MATCAP1 in cell lysates. Scale bar, 5 µm. **(C,D)** Simultaneous visualization of ΔY- and ΔC2-MT biogenesis. (**C**) Representative kymographs showing ΔY-Fab^488^ (green) and ΔC2-Fab^647^ probe (magenta) labeling of the same microtubule over time after the addition of 1 nM Halo-MATCAP1 in cell lysate. Time is shown on the y-axis (scale bar, 1 min), and distance along the microtubule is on the x-axis (scale bar, 5 µm). **(D)** Quantification of the mean fluorescence intensity of ΔY-Fab^488^ (green) and ΔC2-Fab^647^ probes (magenta) along microtubules over time. Data are presented as mean ± S.D., with n = 29 microtubules from three independent experiments.

A potential concern with these assays stems from the fact that both Fab probes recognize the CTT of α-tubulin, which could lead to competition between the probes for substrate binding. To test whether the ΔY-Fab^488^ probe obstructs ΔC2-Fab^647^ binding, we compared ΔC2-microtubule biogenesis by MATCAP1 in the absence and presence of the ΔY-Fab^488^ probe. Our results show that the ΔY-Fab^488^ probe does not affect the kinetics of ΔC2-Fab^647^ microtubule labeling in the presence of MATCAP1 (**Figure S3, A and B**), indicating that the ΔY-Fab^488^ does not hinder binding of the ΔC2-Fab^647^. To provide a molecular explanation, we examined the binding kinetics of the probes. To do this, we treated Taxol-stabilized HeLa microtubules with VASH1/SVBP or MATCAP1 enzymes to generate ΔY-microtubules or ΔC2-microtubules, respectively. The enzymes were washed away by high-salt buffer (**Figure S4**) and then the ΔY-Fab^488^ or ΔC2-Fab^647^ probes were added to the flow chamber. On microtubules with low levels of modification, the ΔY-Fab^488^ and ΔC2-Fab^647^ probes showed rapid binding and unbinding events (**Figure S3, C and D**). On highly modified microtubules (high enzyme amount or long incubation time), the probes appeared to stably decorate microtubules (**Figure S3, E and F**) which likely reflects the binding of multiple probes to α-tubulin CTTs within each pixel. On highly modified microtubules, both the ΔY-Fab^488^ and ΔC2-Fab^647^ probes showed no delay in microtubule binding (**Figure S3, E and F**). Together, these results suggest that ΔY-Fab^488^ and ΔC2-Fab^647^ probes behave similarly and can be used for real-time visualization of microtubule modification.

### MATCAP1 binding and activity are not affected by the detyrosination state of the microtubule

To understand how the state of the microtubule lattice impacts MATCAP1’s enzymatic activity, we characterized MATCAP1 binding to microtubules with different tubulin states using a microscopy-based microtubule binding assay (**Figure 4A**). Halo-MATCAP1 was expressed in COS-7 cells and labeled with the JFX554 Halo ligand. The resulting cell extracts were added to Taxol-stabilized HeLa microtubules to observe the behavior of individual Halo^554^-MATCAP1 molecules.

**Figure 4.**
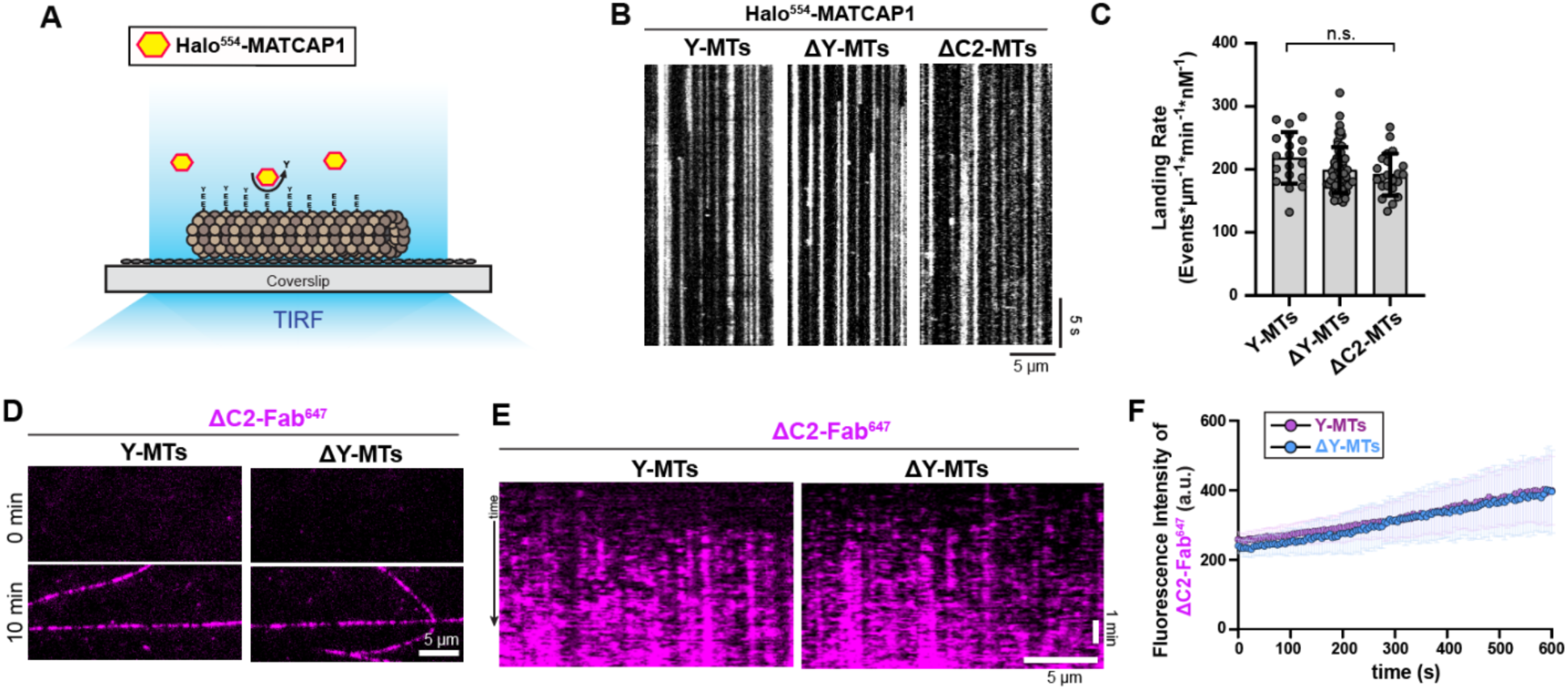
MATCAP1 binding and activity are not affected by the microtubule detyrosination state. **(A)** Schematic of the single-molecule microtubule binding assay. Microtubules were incubated with fluorescently-labeled MATCAP1 and binding events were monitored over time using TIRF microscopy. **(B,C)** Landing rate of MATCAP1 on microtubules with different a-tubulin CTT modifications. Taxol-stabilized HeLa microtubules were untreated (Y-MTs) or were incubated with cell lysates containing VASH1/SVBP to generate ΔY-MTs or with MATCAP1 to generate ΔC2-MTs. The enzymes were removed with a high salt wash and then cell lysate containing Halo^554^-MATCAP1 was added to the flow chamber. (**B**) Representative kymographs showing single molecules of 10 pM Halo^554^-MATCAP1 binding to the indicated microtubules. Time is shown on the y-axis (scale bar, 5 sec) and distance along the microtubule is on the x-axis (scale bar, 5 µm). **(C)** Quantification of the landing rate of Halo^554^-MATCAP1 along microtubules. Each point represents the landing rate of MATCAP1 on an individual microtubule. Data are presented as mean ± S.D., with n = 19–60 microtubules from two or three independent experiments. n.s., not significant (one-way ANOVA). **(D-F)** MATCAP1 activity along Y-MTs and ΔY-MTs. Taxol-stabilized HeLa microtubules were untreated or were incubated with cell lysates containing VASH1/SVBP to generate ΔY-MTs. The enzymes were removed with a high salt wash and then MATCAP1 and the ΔC2-Fab^647^ probe were added to the flow chamber. (**D**) Representative images of ΔC2-Fab^647^ labeling of microtubules at 0 min and after 10 min incubation with 1 nM Halo-MATCAP1. Scale bar, 5 µm. (**E**) Representative kymographs showing ΔC2-Fab^647^ probe labeling over time. Time is shown on the y-axis (scale bar, 1 min), and distance along the microtubule is on the x-axis (scale bar, 5 µm). **(F)** Quantification of the mean fluorescence intensity of ΔC2-Fab^647^ probe for the microtubules in (E). Data are presented as mean ± S.D., with n = 17–23 microtubules from two independent experiments.

We first examined how modification of the α-tubulin CTT affects the binding of MATCAP1 to microtubules. Taxol-stabilized HeLa microtubules were untreated to maintain the tyrosinated state of α-tubulin or were treated with cell lysate containing VASH1/SVBP to generate ΔY-microtubules or were treated with cell lysate containing MATCAP1 to generate ΔC2-microtubules. After washing out the enzymes with high-salt buffer, Halo^554^-MATCAP1 was added to the flow chamber and imaged over time. When added at 10 pM, individual Halo^554^-MATCAP1 molecules were found to bind statically to Y-microtubules, ΔY-microtubules, and ΔC2-microtubules with comparable landing rates (**Figure 4, B and C**). These results also indicate that once bound to Taxol-stabilized microtubules, Halo^554^-MATCAP1 rarely detaches, exhibiting a long dwell time on microtubules (>30 s, the duration of our imaging period). This is in contrast to VASH1/SVBP, which rapidly binds and unbinds from microtubules [dwell time ∼1 s (Yue 2023, Ramirez-Ramos 2022)].

To examine whether prior detyrosination affects the activity of MATCAP1, we used the ΔC2-Fab^647^ probe. Taxol-stabilized HeLa microtubules were untreated or were treated with VAH1/SVBP enzyme to generate ΔY-microtubules. After a high-salt wash to remove VASH1/SVBP, cell lysate containing MATCAP1 was added together with the ΔC2-Fab^647^ probe. No differences were observed for ΔC2-Fab^647^ probe labeling of Y-microtubules versus ΔY-microtubules (**Figure 4, D-F**), suggesting comparable enzymatic activity of MATCAP1 in generating ΔC2-microtubules from both substrates. Overall, these results demonstrate that MATCAP1 binding and activity are not affected by the detyrosination state of the α-tubulin CTT.

### Microtubule binding of MATCAP1 depends on the conformational state of tubulin

We next examined how the conformational state of tubulin in the microtubule lattice affects the binding and activity of MATCAP1. HeLa tubulin was polymerized in the presence of the nonhydrolyzable GTP analog GMPCPP (GMPCPP-microtubules) or in the presence of GTP to generate GDP-microtubules as GTP is converted to GDP upon polymerization. To delay disassembly of GDP-microtubules, glycerol was added during the assay. GMPCPP-microtubules were also maintained in the same buffer. Individual Halo^554^-MATCAP1 molecules exhibited long dwell times on both types of microtubules (**Figure 5A**), however, the landing rate was higher on GMPCPP-microtubules than on GDP-microtubules (**Figure 5B**). These results demonstrate that microtubule binding of MATCAP1 is regulated by the nucleotide state of tubulin within the microtubule lattice. This behavior is in contrast to that of VASH1/SVBP which displays similar landing rates on GMPCPP- and GDP-microtubules (Yue et al. 2023).

**Figure 5.**
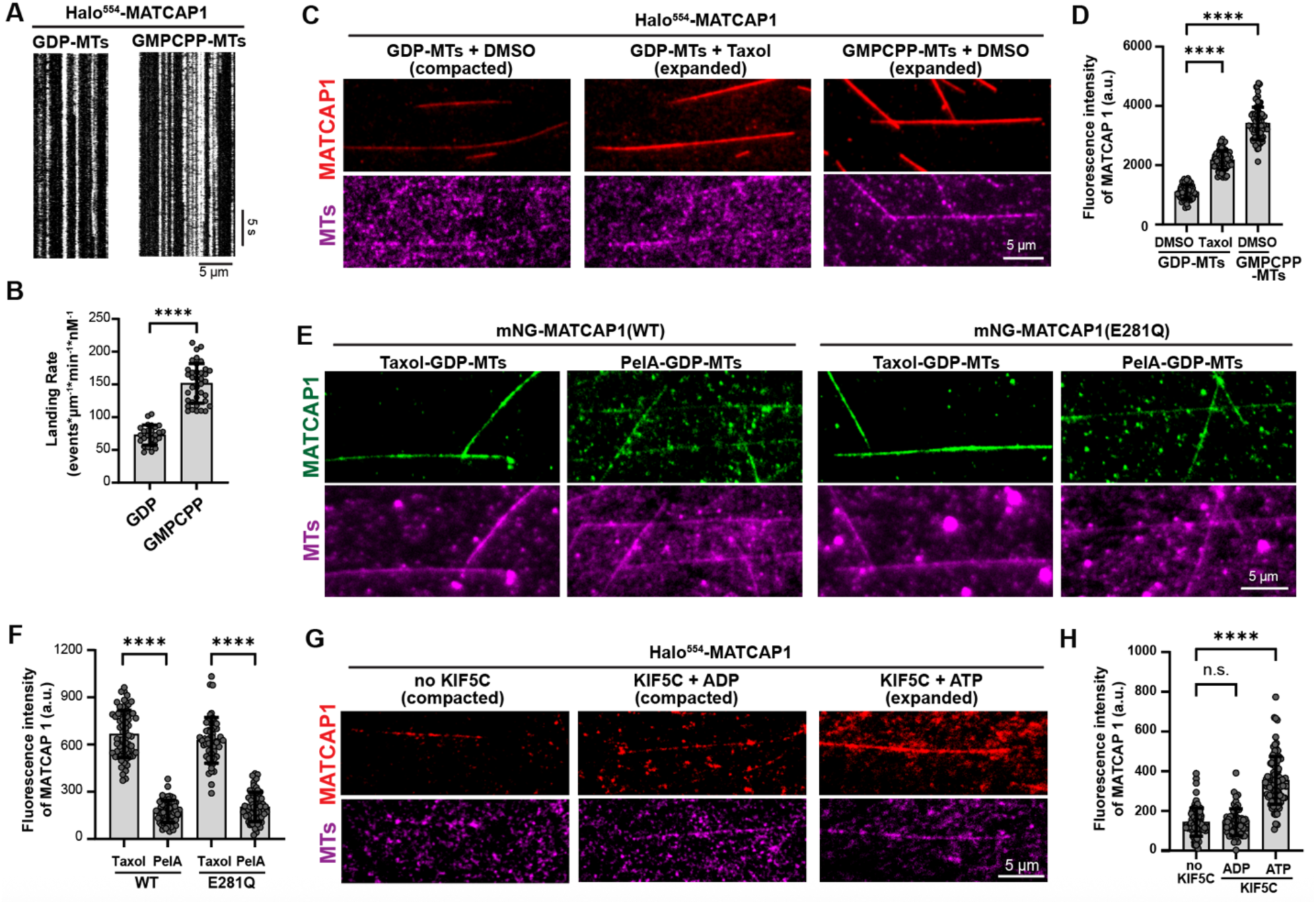
Microtubule binding of MATCAP1 is regulated by the conformational state of tubulin within the microtubule lattice. **(A,B)** Landing rate of Halo^554^-MATCAP1 on GDP-MTs vs GMPCPP-MTs. (**A**) Representative kymographs showing single molecules of 10 pM Halo^554^-MATCAP1 in cell lysates binding to glycerol-stabilized GDP-MTs or GMPCPP-MTs. Time is on the y-axis (scale bar, 5 sec), and distance along the microtubule is on the x-axis (scale bar, 5 µm). **(B)** Quantification of the landing rate of Halo^554^-MATCAP1 along the microtubules in (A). Each point represents the landing rate of MATCAP1 on an individual microtubule. Data are presented as mean ± S.D., with n = 26–37 microtubules from two independent experiments. ****p<0.001 (two-tailed, *t*-test). **(C,D)** Steady-state binding of MATCAP1 to compacted vs expanded microtubules. **(C)** Representative images of 1 nM Halo^554^-MATCAP1 in cell lysates binding to glycerol-stabilized GDP-MTs with DMSO, GDP-MTs with Taxol, or GMPCPP-MTs with DMSO. Scale bar, 5 µm. **(D)** Quantification of the mean fluorescence intensity of Halo^554^-MATCAP1 along microtubules. Data are presented as mean ± S.D., with n = 56–83 microtubules from three independent experiments. ****p<0.0001 (two-tailed, *t*-test). **(E,F)** Steady-state binding of MATCAP1 to Taxol-stabilized vs PelA-stabilized GDP-MTs. **(E)** Representative images of 2 nM WT or E281Q mNG-MATCAP1 in cell lysates binding to Taxol- or PelA-stabilized GDP-MTs. Scale bar, 5 µm. **(F)** Quantification of mean fluorescence intensity of WT and E281Q mNG-MATCAP1 along microtubules. Data are presented as mean ± S.D. with n = 51-72 microtubules from two independent experiments. ****p<0.0001 (two-tailed, *t*-test). **(G,H)** Steady-state binding of MATCAP1 to KIF5C-expanded microtubules. **(G)** Representative images of 3 nM purified Halo^554^-MATCAP1 protein binding to glycerol-stabilized GDP-MTs in the absence of KIF5C (microtubule compacted), in the presence of 100 nM KIF5C and 2 mM ADP (KIF5C inactive, microtubule compacted) or in the presence of 100 nM KIF5C and 2 mM ATP (KIF5C active, microtubule expanded). **(H)** Quantification of the mean fluorescence intensity of Halo^554^-MATCAP1 along microtubules. Data are presented as mean ± S.D., with n = 60–100 microtubules from two or three independent experiments. ****p<0.0001 and n.s., not significant (two-tailed, *t*-test).

The higher landing rate on GMPCPP-microtubules suggests that MATCAP1 may preferentially bind to tubulin within the microtubule lattice that is in an expanded state. To test this, we took advantage of the fact that Taxol can revert GDP-tubulin to an expanded state within the microtubule lattice (Alushin et al. 2014, Prota et al. 2023). We compared the fluorescence intensity of Halo^554^-MATCAP1 along GDP-microtubules with DMSO (compacted state), GDP-microtubules with Taxol (expanded state), and GMPCPP-microtubules with DMSO (expanded state). Halo^554^-MATCAP1 showed 2.1-fold higher binding to Taxol-treated GDP-microtubules than to GDP-microtubules and 3.4-fold higher binding to GMPCPP-microtubules than to GDP-microtubules (**Figure 5, C and D**). To confirm that the preferential binding of MATCAP1 to expanded microtubules was not an artifact of the cell lysate system, we added a dual Strep tag to Halo-MATCAP1 (TwinStrep-Halo-MATCAP1) and purified the protein from COS-7 cells using affinity chromatography (**Figure S5A**). The purified TwinStrep-Halo^554^-MATCAP1 protein showed 2.4-fold higher binding to Taxol-treated GDP-microtubules than to GDP-microtubules and 3.9-fold higher binding to GMPCPP-microtubules than to GDP-microtubules (**Figure S5, B and C**).

As a second test of whether MATCAP1 preferentially binds to tubulins in an expanded state, we compared its ability to bind to GDP-microtubules stabilized with Taxol vs peloruside A (hereafter referred to as PelA). Although both natural products stabilize GDP-microtubules, only Taxol drives tubulin to an expanded state in the microtubule lattice (Huzil et al. 2008, Estevez-Gallego et al. 2023). For these experiments, we used MATCAP1 labeled with mNeonGreen tag at the N-terminus (mNG-MATCAP1) to rule out any influence of the fluorescent tag on MATCAP1 activity. We also examined the microtubule-binding behavior of a catalytically inactive MATCAP1 mutant, E281Q, which localizes to microtubules in cells (Landskron et al. 2022). GDP-microtubules were polymerized from HeLa tubulin, stabilized with either Taxol or PelA, and then incubated with mNG-MATCAP1 or mNG-MATCAP1(E281Q) in an imaging chamber. The fluorescence intensity of mNG-MATCAP1 was 3.8-fold higher on Taxol-GDP-microtubules compared to PelA-GDP-microtubules (**Figure 5, E and F**). Similarly, the E281Q mutant protein showed 3.1-fold higher fluorescence intensity on Taxol-GDP-microtubules than on PelA-GDP-microtubules (**Figure 5, E and F**). These results demonstrate that both WT and E281Q MATCAP1 proteins display preferential binding to tubulin in an expanded state within the microtubule lattice.

Finally, we used the kinesin-1 motor KIF5C as an alternative, non-drug-based mechanism to generate expanded tubulin within the microtubule lattice. Previous work has shown that kinesin-1 triggers lattice expansion when it is in the ATP-bound state but not in the ADP-bound state (Peet et al. 2018, Shima et al. 2018). We purified KIF5C(1-560)-Halo-TwinStrep from insect cells and TwinStrep-Halo^554^-MATCAP1 from COS7 cells and compared MATCAP1 binding to GDP-microtubules in the absence of KIF5C and in the presence of KIF5C under both ATP and ADP conditions. Our results revealed that the presence of active KIF5C (ATP-bound) increased the mean fluorescence intensity of Halo^554^-MATCAP1 along the microtubule lattice by 2.5-fold compared to the absence of KIF5C, whereas KIF5C in the ADP-bound state did not affect MATCAP1 binding (**Figure 5, G and H**). Collectively, these results demonstrate that the microtubule binding of MATCAP1 is regulated by the conformational state of tubulin within the microtubule lattice. This behavior contrasts with that of VASH1/SVBP, which binds equally to microtubules regardless of tubulin conformation (Yue et al. 2023).

### Preferential binding to expanded tubulin determines the ability of MATCAP1 to detyrosinate tubulin in the microtubule lattice

Given the preferential binding of MATCAP1 to an expanded microtubule lattice, we examined whether its detyrosination activity also depends on the conformational state of tubulin in the microtubule lattice. Halo-MATCAP1 in cell lysate was incubated for 6 min with GDP-microtubules with DMSO (compacted state), GDP-microtubules with Taxol (expanded state), or GMPCPP-microtubules with DMSO (expanded state). Bound MATCAP1 was then removed by washing with high-salt buffer. Microtubules were subsequently incubated with the ΔY-Fab^488^ probe to visualize the extent of detyrosination catalyzed by MATCAP1 (**Figure 6A**). We found that MATCAP1 exhibited higher detyrosination activity on GMPCPP-microtubules and Taxol-treated GDP-microtubules than on GDP-microtubules (**Figure 6, B and C**). To ensure that the cell lysate did not affect MATCAP1 activity, we carried out similar experiments with purified TwinStrep-Halo-MATCAP1 protein and found a similar increase in MATCAP1 activity along microtubules with an expanded lattice (**Figure S5, D and E**). These results suggest that the detyrosination activity of MATCAP1 depends on the conformational state of tubulin within the microtubule lattice.

**Figure 6.**
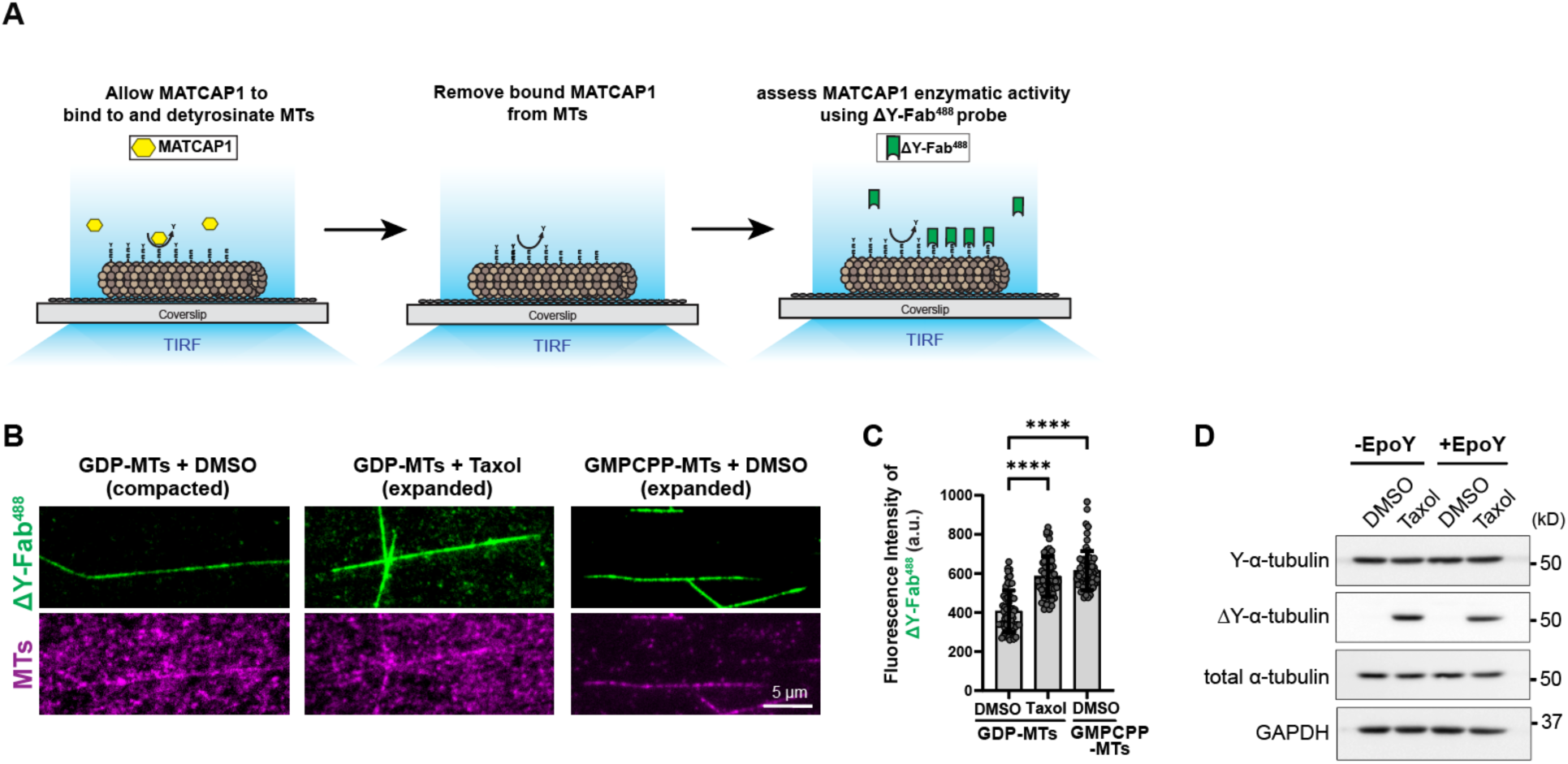
Detyrosination activity of MATCAP1 is regulated by the conformational state of tubulin within the microtubule lattice. **(A)** Schematic of the modified microscopy-based enzymatic assay. Glycerol-stabilized HeLa microtubules were incubated with MATCAP1 (yellow) in cell lysate for microtubule detyrosination. The enzyme was removed by high-salt wash buffer and then the ΔY-Fab^488^ probe was added to visualize microtubule detyrosination. **(B)** Representative images of ΔY-Fab^488^ labeling of glycerol-stabilized GDP-MTs with DMSO, GDP-MTs with Taxol, or GMPCPP-MTs with DMSO after incubation with 1 nM Halo-MATCAP1. Scale bar, 5 µm. **(C)** Quantification of the mean fluorescence intensity of ΔY-Fab^488^ probe labeling along the microtubules in (B). Each point represents the mean fluorescence intensity of ΔY-Fab^488^ along an individual microtubule. Data are presented as mean ± S.D., with n = 61–65 microtubules from three independent experiments. ****p<0.0001 (two-tailed, *t*-test). **(D)** Western blot showing the extent of ΔY modification of microtubules from HeLa cells treated with DMSO or Taxol, in the absence or presence of EpoY, a VASH1/SVBP inhibitor.

As an alternative method to test whether MATCAP1 activity is influenced by the expanded state of the microtubule lattice, we treated HeLa cells with 10 µM Taxol in the absence or presence of EpoY, a suicide inhibitor of VASH1/SVBP (Aillaud et al. 2017). We reasoned that when EpoY blocks the activity of VASH1/SVBP, any microtubule detyrosination activity in the presence of this compound must be due to MATCAP1. We observed that Taxol stimulates microtubule detyrosination in both the absence and presence of EpoY (**Figure 6D**), suggesting that the expanded state of the microtubule lattice is a key feature that gates microtubule detyrosination by MATCAP1.

## Discussion

How cells select specific microtubules for tubulin PTMs remains a major unresolved question in the field. In this study, we provide mechanistic insights into the microtubule binding and enzymatic activity of MATCAP1. By developing microscopy-based enzymatic assays with PTM-specific Fab probes, we discovered that MATCAP1 binds preferentially to microtubules with an expanded lattice to carry out the sequential generation of ΔY- and ΔC2-α-tubulin modifications. Our findings provide insights into how the conformational state of tubulin subunits within the microtubule lattice participate in the spatiotemporal regulation of tubulin PTMs.

A significant challenge in studying tubulin PTMs has been the lack of tools for real-time detection, limiting our ability to understand how specific PTMs are dynamically established and regulated in cells. To address this, we previously developed a fluorescently-labeled Fab probe for ΔY-microtubules (Yue et al. 2023). In this study, we applied the same approach to generate a fluorescently-labeled Fab probe specific for ΔC2-microtubules. Both Fab probes exhibit immediate and selective binding to their respective targets, enabling real-time visualization of ΔY and ΔC2 modifications on individual microtubules. Furthermore, both Fab probes exhibit rapid and transient binding kinetics and thus do not interfere with each other’s labeling under experimental conditions, enabling the simultaneous tracking of both PTMs. Using the dual-Fab strategy, we were able to directly demonstrate that MATCAP1 catalyzes the formation of ΔY-tubulin faster than ΔC2-tubulin on the same microtubules. Collectively, our work establishes Fab-based probes as powerful tools for spatiotemporal analysis of microtubule PTMs, opening new avenues to investigate the functional crosstalk among distinct tubulin PTMs.

We demonstrate that MATCAP1 efficiently generates both ΔY- and ΔC2-microtubules in vitro and in cells, consistent with recent work (Nicot et al. 2023). However, it had remained unclear whether MATCAP1 acts independently of CCP1 to generate ΔC2-microtubules in cells. Here, by overexpressing MATCAP1 in Δ*TTL*Δ*CPP1* HeLa cells, we demonstrate that MATCAP1 is capable of directly generating ΔC2-α-tubulin. This cell-based evidence is corroborated by our in vitro enzymatic assays, which reveal that MATCAP1 efficiently generates the ΔC2 modification along microtubules.

The ability of MATCAP1 to generate both ΔY- and ΔC2-α-tubulin modifications may be facilitated by its relatively long dwell time (>30 s) on microtubules. In this respect, MATCAP1 behavior is more similar to VASH2/SVBP (dwell time >11 s) than to VASH1/SVBP (dwell time ∼1 s) (Ramirez-Rios et al. 2023, Yue et al. 2023). It seems possible that MATCAP1 could remain bound to a single tubulin substrate and sequentially modify its αCTT. This possibility is supported by structural studies showing that MATCAP1 interacts with αCTT residues N-terminal to the cleavage site and is able to carry out a variety of cleavage events at the C-terminal end of the αCTT (Landskron et al. 2022). However, we cannot exclude the possibility that individual MATCAP1 proteins diffuse with a diffraction-limited area to proteolyze the CTTs of multiple tubulins before falling off of the microtubule.

We find that the microtubule binding and enzymatic activity of MATCAP1 are insensitive to the prior detyrosination state of tubulin in the microtubule lattice, similar to VASH1/SVBP and VASH2/SVBP (Ramirez-Rios et al. 2023, Yue et al. 2023). These findings rule out a model in which progressive modification occurs through a positive feedback mechanism caused by increased affinity of the enzyme for its modified substrate. Rather, we find that MATCAP1 binds preferentially to microtubules with an expanded lattice conformation, which can be induced by preventing β-tubulin GTP hydrolysis, Taxol treatment, or stepping of active kinesin-1 along the microtubule. This binding behavior is different than that of VASH1/SVBP which binds equally to microtubules that are in a compacted vs expanded state (Yue et al. 2023). The molecular basis for this difference is not readily apparent from recent cryo-EM data. VASH1/SVBP binds in between protofilaments and engages the two adjacent α-tubulin subunits, whereas MATCAP1 binds along one protofilament of a microtubule, primarily contacting the α-tubulin subunit targeted for cleavage while making contacts with an adjacent α-tubulin subunit (Li et al. 2020, Landskron et al. 2022, Ramirez-Rios et al. 2023). Thus, both MATCAP1 and VASH1/SVBP make extensive contacts with α-tubulin beyond its CTT, yet only MATCP1 shows increased binding to α-tubulin in an expanded state. Further structural studies are needed to understand the interactions of full-length MATCAP1 and VASH1/SVBP proteins along microtubules in different conformational states.

Our findings that the detyrosination activities of MATCAP1 (this study) and VASH1/SVBP (Yue et al. 2023) are regulated by the conformational state of tubulin within the microtubule lattice are consistent with emerging studies showing that tubulin conformation can influence the binding affinity and activity of various MAPs and vice versa. For example, kinesin-1, CAMSAP3, and MAP7 preferentially bind to expanded microtubules, and their binding can further promote lattice expansion (Peet et al. 2018, Shima et al. 2018, Liu and Shima 2023, Shen and Ori-McKenney 2024). In contrast, Doublecortin (DCX), MAP2, and tau, preferentially bind to and induce lattice compaction (Castle et al. 2020, Siahaan et al. 2022, Paquette et al. 2025). Collectively, our findings support a model in which MAPs that modulate tubulin conformation within the microtubule lattice contribute to the regulation of tubulin PTMs.

## Materials and Methods

### Plasmids

For the Halo-MATCAP1 construct, a gene fragment containing Halo and human MATCAP1 (amino acids 1-471, UniProt Q68EN5) was synthesized and inserted into the pC1 vector using the NEBuilder HiFi DNA assembly kit. Additional MATCAP1 constructs were generated by replacing the Halo tag with monomeric Neon Green (mNG-MATCAP1), tandem PA (GVAMPGAEDDVV) and TagRFP tags (PA-TagRFP-MATCAP1), or tandem TwinStrep [WSHPQFEKGGGSGGGSGGSAWSHPQFEK (Schmidt et al. 2013)] and Halo tags (TwinStrep-Halo-MATCAP1) via PCR and Gibson assembly (NEB HiFi kit). The E281Q mNG-MATCAP1 mutant was generated by site-directed mutagenesis. To generate a construct for stable knock-in of Halo-MATCAP1 in HeLa Kyoto cells, the Halo-MATCAP1 fragment was PCR amplified and inserted into pEM791 vector that had been digested with BsrGI and AgeI. The resulting pEM791-Halo-MATCAP1 vector was used to establish knock-in HeLa cell lines expressing Halo-MATCAP1 in a doxycycline inducible manner using recombination mediated cassette exchange (Khandelia et al. 2011). A truncated, constitutively active rat kinesin-1 [KIF5C(1-560)] was amplified by PCR and subcloned into the pFastBac1 vector to generate KIF5C(1-560)]-Halo-TwinStrep tag for protein expression and purification. All plasmids were verified by DNA sequencing.

### Cell culture, transfection, immunofluorescence, and lysate preparation

HeLa Kyoto cells (RRID: CVCL_1922), Δ*TTL*Δ*CCP1* HeLa cells (Hotta et al. 2023), and stable Halo-MATCAP1 HeLa cells were cultured in Gibco DMEM high glucose (Invitrogen 11965092) with 10% tetracycline-negative FBS (R&D Systems, Cat# S10350), 1% GlutaMAX (Invitrogen 35050061), and 1% PenicillinStreptomycin (10000 UmL) (Invitrogen 15140163) and grown at 37°C with 5% CO_2_.

HeLa cells seeded on coverslips in a 6-well plate were transfected using Lipofectamine 2000 (Invitrogen, Cat# 11668030) and used the following day. For the Halo-MATCAP1 stable HeLa cell line, Halo-MATCAP1 expression was induced by adding 2 µg/mL doxycycline to the culture medium along with Halo554 ligand (to label the protein) 16 hours prior to fixation.

Cells were fixed with ice-cold methanol for 10 min at −20°C. Blocking was performed with 2% BSA in TBS (0.2 M Tris Base, 137 mM NaCl, pH 7.6) supplemented with 0.1% Triton X-100. Antibodies used were: anti-ΔY-α-tubulin antibody (Clone RM444; RevMAb Biosciences, Cat#31-1335-00, final concentration 1 mg/ml, 1 hour), anti-ΔC2-α-tubulin antibody (Clone RM447; RevMAb Biosciences, Cat# 31-1339-00, final concentration 1 mg/ml, 1 hour), anti-PA-tag antibody (clone NZ-1, Fujifilm Wako Pure Chemical, Cat# 016-25861, diluted at 1:250, 45 min), DM1a-AlexaFluor647 (for PA-TagRFP-MATCAP1 experiment) or FITC-conjugated DM1a (for Halo-MATCAP1 experiment; Sigma-Aldrich, Cat# F2168, 1:500). All antibodies were diluted in the blocking solution. DNA was stained with 5 mg/mL Hoechst. Coverslips were mounted with ProLong Diamond (Thermo Fisher). Images were obtained with a DeltaVision microscope equipped with an Olympus Plan Apo N 60x/1.42 oil immersion lens. Images were deconvolved, and single optical sections are presented.

COS-7 [male *Ceropithecus aethiops* (green monkey) kidney fibroblast, RRID: CVCL_0224] cells were grown in Dulbecco’s modified Eagle medium (DMEM, Gibco 11960044) supplemented with 10% (vol/vol) Fetal Clone III (HyClone SH3010903) and 2 mM GlutaMAX (L-alanyl-L-glutamine dipeptide in 0.85% NaCl, Gibco 35050061) and grown at 37°C with 5% (vol/vol) CO_2_.

COS-7 cells were transfected using Trans-IT LT1 (Mirus) and Opti-MEM (Thermo Fisher) according to the manufacturer’s instructions. Halo-tagged protein was fluorescently labeled by adding 50 nM JFX554 Halo ligand (Tocris Bioscience) to the cell medium after the transfection. The cells were harvested 16h post-transfection by low-speed centrifugation at 1,500 g for 5 min at 4°C. The pellet was rinsed once in PBS and resuspended in ice-cold BRB80 buffer (80 mM Pipes/KOH pH 6.8, 1 mM MgCl_2_, and 1 mM EGTA) freshly supplemented with 1 mM phenylmethylsulfonylfluoride (PMSF) and protease inhibitors (Sigma-Aldrich). After lysing the cells on ice by a Sonic Dismembrator with a microtip (Fisher Scientific), the lysate was clarified by centrifugation at 20,000 g for 10 min at 4 °C. The supernatant was aliquoted, snap-frozen in liquid nitrogen, and stored at 80 °C until further use.

The concentration of Halo-MATCAP1 in the lysates was measured by a dot-blot, in which 1 µL of COS-7 lysates expressing Halo-MATCAP1 and a series of KIF5C(1-560)-Halo protein of known concentrations were spotted onto a nitrocellulose membrane. The membrane was air-dried for 30 min to 1 h and immunoblotted with a primary antibody to Halo tag (Promega, G9281) at room temperature for 1 h and subsequentially with a secondary antibody 680nm anti-rabbit (Jackson ImmunoResearch Laboratories Inc.) at room temperature for 30 min. The fluorescence intensity of the spots on the nitrocellulose membrane was detected by Azure 600 (Azure Biosystems) and quantified based on the standard curve of known concentration of KIF5C(1-560)- Halo protein using Fiji/ImageJ (NIH).

### Western blot

HeLa cells were lysed in NP-40 buffer (6 mM Na_2_HPO_4_, 4 mM NaH_2_PO_4_, 2 mM EDTA, 150 mM NaCl, 1% NP40 and protease inhibitors) and sonicated. Lysates were clarified centrifugation at 21,000 x g for 15 min at 4°C. Protein concentrations were determined using the Bradford protein assay using BSA as a standard. For SDS-PAGE, 15 µg of total protein per sample was loaded. Standard 10% acrylamide Laemmli gels were used, except for experiments presented in Figure S1, where high pH separation gels were used to resolve α- and β-tubulin as previously described (Banerjee et al. 2010). Proteins were transferred to nitrocellulose membranes, which were then blocked with 5% skim milk in PBS supplemented with 0.05% tween-20 (PBST) for 1 hour at room temperature. Membranes were incubated with primary antibodies at 4°C overnight. Following three washes with PBST, membranes were incubated with fluorescently labeled secondary antibodies for 1 hour at room temperature, followed by another three washes. Fluorescent signals were detected using an Azure 600 imaging system (Azure Biosystems). Antibodies used: anti-tyrosinated α-tubulin rat monoclonal clone YL1/2 (Accurate Chemical and Scientific, Cat# YSRTMCA77G, 1:3,000); anti-detyrosinated α-tubulin rabbit monoclonal RM444 (RevMAb Biosciences, Cat#31-1335-00, 0.1 mg/ml); anti-ΔC2 α-tubulin rabbit monoclonal RM447 (RevMAb Biosciences, Cat# 31-1339-00, 0.5 mg/ml); anti-α-tubulin mouse monoclonal DM1a (Millipore Sigma, Cat# 05-829, 1:3,000), anti-β-tubulin mouse monoclonal E7 (DSHB); anti-PA-tag rat monoclonal NZ-1 (FUJIFILM Wako Pure Chemical, Cat# 016-25861, 1:5,000); anti-GAPDH mouse monoclonal G-9 (Santa Cruz, Cat# sc-365062, 1:2,000).

### Protein purification

For KIF5C(1-560)-Halo-TwinStrep protein expression and purification, Sf9 cells were cultured in suspension with serum-free sf900 II SFM medium (Thermo Fisher Scientific) supplemented with Antibiotic-Antimycotic (Gibco) in flasks at 28°C in a non-CO_2_ non-humidified incubator with an orbital shaker platform set at 110 rpm. The cells were infected with baculovirus generated according to the Bac-to-Bac system (Invitrogen). In brief, pFastBac1 plasmid was transformed into DH10Bac *Escherichia coli* to generate recombinant bacmids. Bacmid DNA was isolated with the HiPure Plasmid DNA miniprep kit (Invitrogen) and confirmed by PCR analysis. Recombinant bacmid DNA was transfected into Sf9 cells using Cellfectin II (Invitrogen). 7d after transfection, the supernatant containing P1 baculovirus was collected by centrifugation at 3,000 rpm for 3 min at 4°C. The baculovirus was amplified by successive infection of Sf9 cells to generate P2 and P3 baculoviruses. Baculovirus-containing supernatants were stored at 4°C in the dark. To purify protein, Sf9 cells were infected with 3% P3 baculovirus (vol/vol). 3 d after infection, the cells were harvested by centrifugation for 15 min at 3,000 rpm at 4°C. The pellet was washed once with PBS and resuspended in ice-cold lysis buffer (200 mM NaCl, 4 mM MgCl_2_, 0.5 mM EDTA, 1 mM EGTA, 0.5% igepal, 7% sucrose, and 20 mM imidazole-HCl, pH 7.5) supplemented with 2 mM ATP, 1 mM PMSF, 5 mM DTT, and protease inhibitor cocktail. After 30 min incubation on ice, the lysates were clarified by ultracentrifugation for 20 min at 20,000 rpm in F12-8×50y rotor (Sorvall 3421), and the supernatants were incubated with Strep-Tactin beads (Strep-Tactin XT 4Flow resin, IBA) for 1h at 4°C with rotation. Strep-Tactin beads with bound proteins were collected in a PD-10 column and washed with wash buffer (150 mM KCl, 25 mM imidazole-HCl, pH 7.5, 5 mM MgCl_2_, 1 mM EDTA, and 1 mM EGTA) supplemented with 1 mM PMSF, 3 mM DTT, 3 mM ATP, and protease inhibitor cocktail. Bound proteins were eluted with elution buffer (25 mM KCl, 25 mM imidazole-HCl, pH 7.5, 5 mM EGTA, 2 mM MgCl_2_, 2 mM DTT, 0.1 mM ATP, 1 mM PMSF, protease inhibitor cocktail and 10% glycerol) supplemented with 50 mM biotin in 6×0.5 mL fractions. The eluted fractions were analyzed by SDS-PAGE and fractions containing the peak protein were selected by visual inspection of the gel. Protein fractions were combined and dialyzed against dialysis buffer (25 mM imidazole-HCl, pH 7.5, 25 mM KCl, 5 mM EGTA, 2 mM MgCl_2_, 2 mM DTT, 0.1 mM ATP and 10% glycerol). After 2 hrs, the buffer was changed to fresh dialysis buffer and dialyzed overnight at 4°C to remove biotin from the sample. The protein sample was collected by centrifugation, and aliquots were snap frozen in liquid nitrogen and stored in −80°C until further use.

For purification of MATCAP1 protein from COS7 cells, COS-7 cells were transfected with a plasmid for expression of TwinStrep-Halo-MATCAP1 protein and were harvested by centrifugation 16-20 h later. The cells from four 10 cm dishes were lysed in 1 mL of lysis buffer (25 mM HEPES, 115 mM KOAc, 5 mM NaOAc, 5 mM MgCl2, 0.5 mM EGTA, 10% Tx-100, pH 7.4) supplemented with 1 mM PMSF and protease inhibitor cocktail and cleared by centrifugation. The cell lysates were incubated with 100 µl of Strep-Tactin XT 4Flow beads (IBA) for 1h at 4 °C. Beads were washed three times with washing buffer (20 mM HEPES/ KOH pH 7.5, 150 mM NaCl, 1 mM DTT). The proteins were eluted in 1 mL of washing buffer containing 50 mM biotin (Sigma) for 1h at 4 °C. The eluted protein was further concentrated to 100 µL using a10K MWCO spin column (Cytiva). Purified proteins were snap-frozen in liquid nitrogen and stored in - 80°C.

### Preparation of ΔY-Fab^488^ and ΔC2-Fab^647^ probes

Fluorescently-labeled ΔY-Fab^488^ and ΔC2-Fab^647^ probes were generated from rabbit recombinant monoclonal antibodies against ΔY-and ΔC2-α-tubulin (RevMAb Biosceinces, Cat#31-1335-00 and Cat# 31-1339-00, respectively). Fluorescent labeling of the parental IgG and subsequent Fab preparation were prepared using the Fluorescent Protein Labeling Kit (Thermo Fisher, Cat# A10235) and the Fab Preparation Kit (Thermo Fisher, Cat# 44985) according to the manufacturer’s instructions.

### Microtubule polymerization

Tubulin was purified from HeLa cells using TOG affinity chromatography and labeled with fluorescent dyes as described (Thomas et al). For fluorescently-labeled microtubules, the tubulin mix contained 10% Alexa647-labeled or Alexa568-labeled HeLa tubulin. All microtubules were stored in the dark at 37°C until further use.

Taxol-stabilized GDP-microtubules: Microtubules were polymerized from 30 μM HeLa tubulin in BRB80 buffer supplemented with 2.5 mM GTP and 4 mM MgCl_2_ at 37 °C for 35 min. A 5× volume of prewarmed BRB80 buffer containing 10 μM Taxol was added. The microtubules were centrifuged at 15,000 rpm for 10 min at room temperature. The pellet was resuspended in the same 5× volume of prewarmed BRB80 buffer containing 10 μM Taxol.

Taxol-stabilized ΔY-microtubules: Taxol-stabilized HeLa microtubules were incubated with cell lysate containing 0.07–1 nM unlabeled VASH1/SVBP and 10 μM Taxol in P12 buffer (12 mM Pipes/KOH pH 6.8, 1 mM MgCl_2_, 1 mM EGTA) for 5 s - 6 min to generate microtubules with different levels of detyrosination. The enzyme was removed from the microtubules by washing with BRB80 buffer twice.

Taxol-stabilized ΔC2-microtubules: Taxol-stabilized HeLa microtubules were incubated with cell lysate containing 0.7–2 nM MATCAP1 for 3 min - 1 h to generate microtubules with different levels of ΔC2 modification. The enzyme was removed from the microtubules by washing with high-salt buffer (BRB80 buffer supplemented with 200 mM NaCl).

PelA-stabilized GDP-microtubules: Microtubules were polymerized from 30 μM HeLa tubulin in BRB80 buffer supplemented with 2.5 mM GTP and 4 mM MgCl_2_ at 37 °C for 35 min. A 2× volume of prewarmed BRB80 buffer containing 1 μM PelA was added, and the microtubules were centrifuged at 15,000 rpm for 10 min at room temperature. The pellet was resuspended in 3× volume of prewarmed BRB80 buffer containing 1 μM PelA.

GMPCPP-microtubules: Microtubules were polymerized from 5 μM HeLa tubulin in BRB80 buffer supplemented with 4 mM GMPCPP (Jena Bioscience) and 1 mM MgCl_2_ at 37 °C for 35 min. The microtubules were then centrifuged at 15,000 rpm for 10 min at room temperature. The pellet was resuspended in the same volume of prewarmed BRB80 buffer containing 1 mM DTT.

GDP-microtubules: Microtubules were polymerized from 60 µM HeLa tubulin in BRB80 buffer supplemented with 2.5 mM GTP and 5 mM MgCl_2_ at 37 °C for 35 min. A 5× volume of prewarmed BRB80 buffer containing 25% glycerol and 1 mM GTP was added. The microtubules were centrifuged at 15,000 rpm for 10 min at room temperature. The pellet was resuspended in the same 5× volume of BRB80 buffer containing 25% glycerol and 1 mM GTP.

### TIRF microscopy

All *in vitro* assays were performed on an inverted Nikon Ti-E/B total internal reflection fluorescence (TIRF) microscope with a perfect focus system, a 100 × 1.49 NA oil immersion TIRF objective, three 20 mW diode lasers (488 nm, 561 nm, and 640 nm) and EMCCD camera (iXon^+^ DU879; Andor). Image acquisition was controlled using Nikon Elements software. A flow chamber (∼10 μl volume) was assembled by attaching a clean #1.5 coverslip (Fisher Scientific) to a glass slide (Fisher Scientific) with two strips of double-sided tape. All *in vitro* assays were performed in the flow chambers at room temperature.

### Microscopy-based enzymatic assays in vitro

To examine the biogenesis of ΔY- and/or ΔC2-microtubules along Taxol-stabilized HeLa microtubules, polymerized microtubules were introduced into a flow chamber and incubated for 3 min at room temperature to allow for nonspecific adsorption to the coverslip. After washing with blocking buffer (1 mg/mL casein in P12 buffer), the flow chamber was infused with imaging buffer [P12 buffer supplemented with 3 mg/mL casein, 10 µM Taxol, oxygen scavenger mix (1 mM DTT, 1 mM MgCl_2_, 10 mM glucose, 0.2 mg/ml glucose oxidase, and 0.08 mg/ml catalase)] containing VASH1/SVBP-Halo or Halo-MATCAP1( in cell lysate or as purified protein), and 592 pg/μL ΔY-Fab^488^ and/or 5 ng/µL ΔC2-Fab^647^ probe. Images were acquired every 5 s for 10 min or every 30 s for 60 min by TIRF microscopy. For quantification, the mean fluorescence intensity of the ΔY-Fab^488^ and/or ΔC2-Fab^647^ probes along the microtubule in each frame was measured by drawing a line along each microtubule (width= 3 pixels) using Fiji/ImageJ2.

To examine the biogenesis of ΔY-microtubules along HeLa microtubules in different nucleotide or conformational states, GDP-microtubules or GMPCPP-microtubules were introduced into a flow chamber and incubated for 3 min at room temperature to allow for nonspecific adsorption to the coverslip. After washing with blocking buffer (1 mg/mL casein and 10% glycerol in BRB80 buffer), the flow chamber was infused with binding buffer (BRB80 buffer supplemented with 10% glycerol, 0.3 mg/mL casein, Halo-MATCAP1 in cell lysate or purified protein, and 20 µM Taxol or DMSO) and incubated for 6 min at 37 °C to allow MATCAP1 to detyrosinate microtubules. To remove MATCAP1 from the microtubules, the flow chamber was sequentially washed with 1) blocking buffer 2 (1 mg/mL casein and 25% glycerol in BRB80 buffer), 2) blocking buffer 2 supplemented with 200 mM NaCl, and 3) blocking buffer 3 (1 mg/mL casein and 25% glycerol in P12 buffer). Subsequently, the flow chamber was washed with imaging buffer (P12 buffer supplemented with 25% glycerol, 0.3 mg/mL casein, oxygen scavenger mix, and 592 pg/μL ΔY-Fab^488^ probe) and incubated for 5 min. The flow chamber was then sealed with molten paraffin wax and imaged. Snapshots were acquired using a TIRF microscope. The fluorescence intensities along the microtubules were measured using Fiji/ImageJ2 (width= 3 pixels), and the fluorescence intensity of an adjacent region was subtracted to account for background noise. Notably, different buffers and concentrations of glycerol were used at each step to optimize microtubule stability, microtubule binding behavior of MATCAP1, and ΔY-Fab^488^ probe. Our study found that 25% glycerol decreases the microtubule binding of MATCAP1, and BRB80 buffer reduces the labeling of the ΔY-Fab^488^ probe for ΔY-microtubules.

### Microtubule-binding assays in vitro

To examine the landing rate of MATCAP1 on Taxol-stabilized HeLa microtubules, polymerized microtubules were introduced into a flow chamber and incubated for 3 min at room temperature to allow for nonspecific adsorption to the coverslips. After washing with blocking buffer (1 mg/ml casein in P12 buffer), the flow chamber was infused with imaging buffer [P12 buffer supplemented with 10 µM Taxol, 3 mg/ml casein, oxygen scavenger mix, and 10 pM Halo^554^-MATCAP1 in cell lysate]. To examine the landing rate of Halo^JFX554^-MATCAP1 in cell lysate on GDP-microtubules versus GMPCPP-microtubules, 25% glycerol was added to all the buffers to stabilize the microtubules during the assay. The flow chamber was sealed with molten paraffin wax and imaged. Images were acquired continuously every 100 ms for 300 frames by TIRF microscopy. Maximum-intensity projections were generated, and kymographs were produced by drawing along microtubules (width= 3 pixels) using Fiji/ImageJ2. The landing rate was calculated as the number of microtubule binding events per minute per nanomolar protein per micrometer of microtubule.

To examine the microtubule binding of MATCAP1 to HeLa microtubules in different nucleotide or conformational states, GDP-microtubules or GMPCPP-microtubules were added to flow chambers and incubated for 3 min at room temperature to allow for nonspecific adsorption to the coverslips. After washing with blocking buffer (1 mg/mL casein and 10% glycerol in BRB80 buffer), the flow chamber was infused with binding buffer (BRB80 buffer supplemented with 10% glycerol, 0.3 mg/mL casein, oxygen scavenger mix, 1 nM Halo-MATCAP1 in cell lysate or 1.5 nM purified protein, and 20 µM Taxol or DMSO). The flow chamber was then sealed with molten paraffin wax and imaged. Snapshots were acquired using TIRF microscopy. The fluorescence intensities along the microtubules were measured using Fiji/ImageJ2 (width= 3 pixels), and the fluorescence intensity of an adjacent region was subtracted to account for background noise.

For microtubule binding experiments on Taxol- and PelA-stabilized GDP-microtubules, the blocking buffer was 1 mg/ml casein in BRB80 buffer and the imaging buffer was BRB80 buffer supplemented with 1 mg/ml casein, oxygen scavenger mix, and cell lysate containing 2 nM WT or E281Q mNG-MATCAP1.

For microtubule binding experiments during the stepping of mobile KIF5C(1-560) protein, the blocking buffer was 1 mg/mL casein and 25% glycerol in P12 buffer and the imaging buffer was P12 buffer supplemented with 25% glycerol, 0.3 mg/ml casein, oxygen scavenger mix, and 3 nM purified Halo-MATCAP1 protein. The assays were supplemented with purified KIF5C(1-560) protein under three conditions: 1) without KIF5C(1-560)-Halo, 2) with 100 nM KIF5C(1-560)-Halo and 2 mM ATP, or 3) with 100 nM KIF5C(1-560)-Halo, 2 mM ADP, and 2 unit/mL Hexokinase.

## Acknowledgements

We thank Dan Sackett (NIH) for gift of peloruside A. We thank Mike Cianfrocco, Morgan DeSantis, Dave Sept (University of Michigan) and members of their laboratories for advice and feedback.

## Credit

YY - conceptualization, investigation, methodology, formal analysis, validation, data curation, writing - original draft, writing - review and editing

TH - conceptualization, investigation, methodology, formal analysis, validation, data curation, writing - review and editing

RO - conceptualization, funding acquisition, project administration, resources, supervision, writing - original draft, writing - review and editing

KJV - conceptualization, funding acquisition, project administration, resources, supervision, writing - original draft, writing - review and editing

## Abbreviations

ΔY: detyrosinated α-tubulin
ΔC2: α-tubulin with the C-terminal two residues removed
αCTT: C-terminal tail of α-tubulin
PTM: posttranslational modification
MAP: microtubule associated protein
VASH: vasohibin
SVBP: small-vasohibin binding protein
MATCAP: microtubule-associated tyrosine carboxypeptidase

---

**Figure S1.**
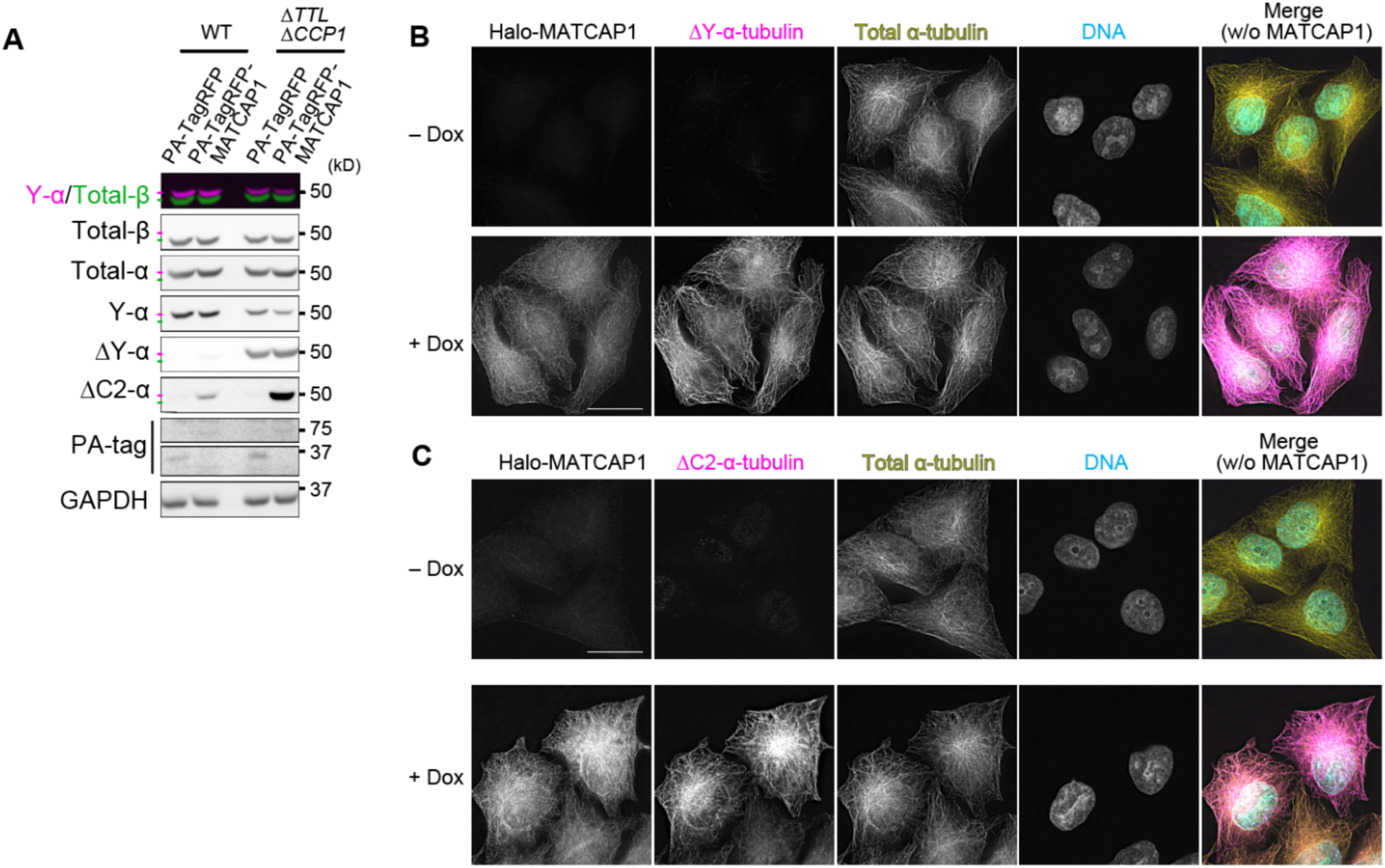
Enzymatic behavior of MATCAP1 in cells. **(A)** Western blot of cell lysates from WT or Δ*TTL*Δ*CCP1* HeLa cells transiently expressing PA-TagRFP or PA-TagRFP-MATCAP1 separated on high pH gels to resolve α- and β-tubulin. Nitrocellular membranes were blotted with antibodies against tyrosinated α-tubulin (Y-α, magenta) and β-tubulin (green) simultaneously or with antibodies against total α-tubulin, ΔY-α-tubulin, ΔC2-α-tubulin, the PA tag, and GAPDH. **(B,C)** Halo-MATCAP1 stable HeLa cells were untreated (-Dox) or treated with doxycycline (+Dox) to induce Halo-MATCAP1 expression. Halo-MATCAP1 was labeled with JFX554 Halo ligand, and then cells were fixed and stained with antibodies against (**B**) ΔY-α-tubulin or (**C**) ΔC2-α-tubulin (magenta) and total α-tubulin (yellow). DNA is shown in cyan. Scale bars, 20 µm.

**Figure S2.**
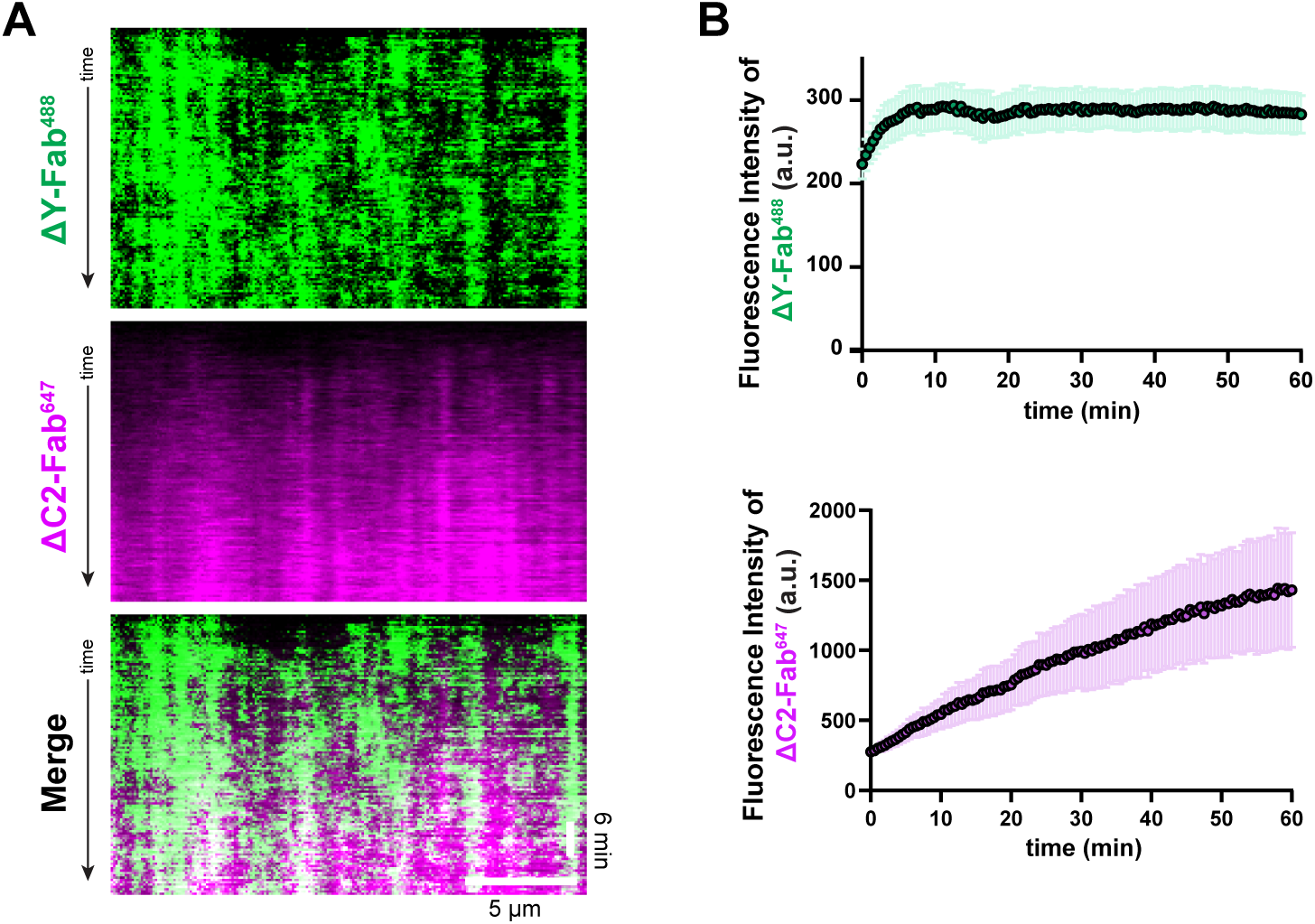
MATCAP1 generates ΔY-MTs faster than ΔC2-MTs in vitro. **(A)** Representative kymographs showing ΔY-Fab^488^ (green) and ΔC2-Fab^647^ (magenta) labeling of Taxol-stabilized HeLa microtubules over 1 h incubation with 4.2 nM Halo-MATCAP1 in cell lysate. Time is shown on the y-axis (scale bar, 6 min), and distance along the microtubule is on the x-axis (scale bar, 5 µm). **(B)** Quantification of the mean fluorescence intensity of ΔY-Fab^488^ (green) and ΔC2-Fab^647^ (magenta) probes along microtubules over time. Data are presented as mean ± S.D., with n = 100 microtubules from two independent experiments.

**Figure S3:**
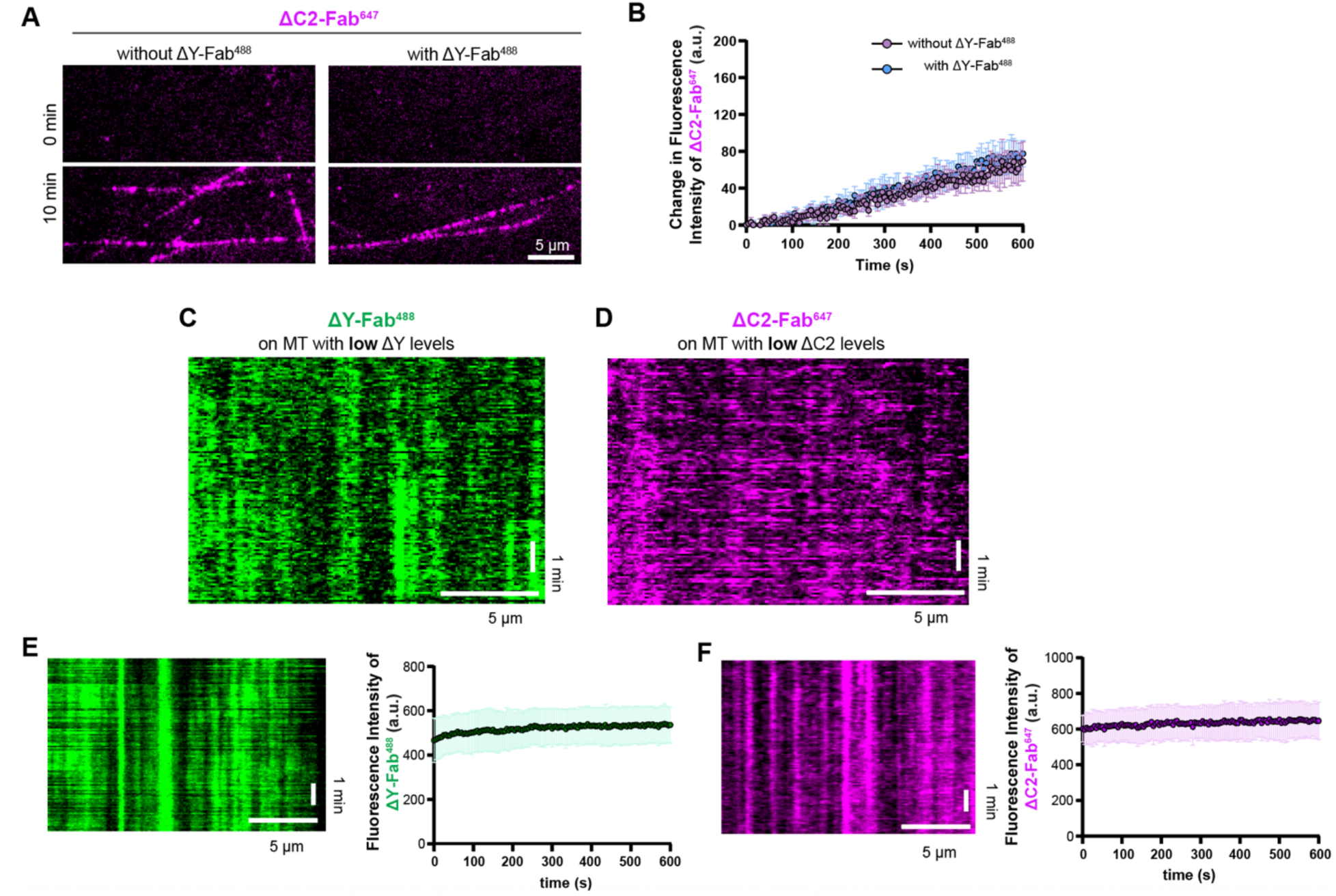
Controls for Fab binding to microtubules. **(A,B)** The ΔY-Fab does not hinder binding of the ΔC2-Fab. **(A)** Representative images of ΔC2-Fab^647^ labeling of Taxol-stabilized HeLa microtubules at 0 min and after 10 min incubation with 1 nM Halo-MATCAP1 in cell lysate, without or with ΔY-Fab^488^. Scale bar, 5 µm. **(B)** Quantification of the mean fluorescence intensity of ΔC2-Fab^647^ probe labeling along microtubules over time. Data are presented as mean ± S.D., with n = 17–23 microtubules from two independent experiments. **(C,D)** Probe binding to microtubules with low levels of modification. Representative kymographs of **(C)** ΔY-Fab^488^ or (**D**) ΔC2-Fab^647^ probe labeling of Taxol-stabilized HeLa microtubules incubated with (C) 0.07 nM VASH1/SVBP for 2–3 s or (D) 0.7 nM MATCAP1 for 3 min to generate low levels of modification. The enzymes were then washed away with high salt buffer and the probes were added to the flow chamber and monitored over time. Time is shown on the y-axis (scale bar, 1 min) and distance along the microtubule is on the x-axis (scale bar, 5 µm). **(E,F)** Probe binding to microtubules with high levels of modification. Left, representative kymographs of ΔY-Fab^488^ or (**F**) ΔC2-Fab^647^ probe labeling of Taxol-stabilized HeLa microtubules incubated with (**E**) 0.7 nM VASH1/SVBP for 2–3 s or (**F**) 1.4 nM MATCAP1 for 15 min. The enzymes were then washed away with high salt buffer and the probes were added to the flow chamber and monitored over time. Time is shown on the y-axis (scale bar, 1 min), and distance along the microtubule is on the x-axis (scale bar, 5 µm). Right, quantification of the mean fluorescence intensity of (E) ΔY-Fab^488^ probe and (F) ΔC2-Fab^647^ along microtubules. Data are presented as mean ± S.D., with n = 13–21 microtubules from two independent experiments.

**Figure S4.**
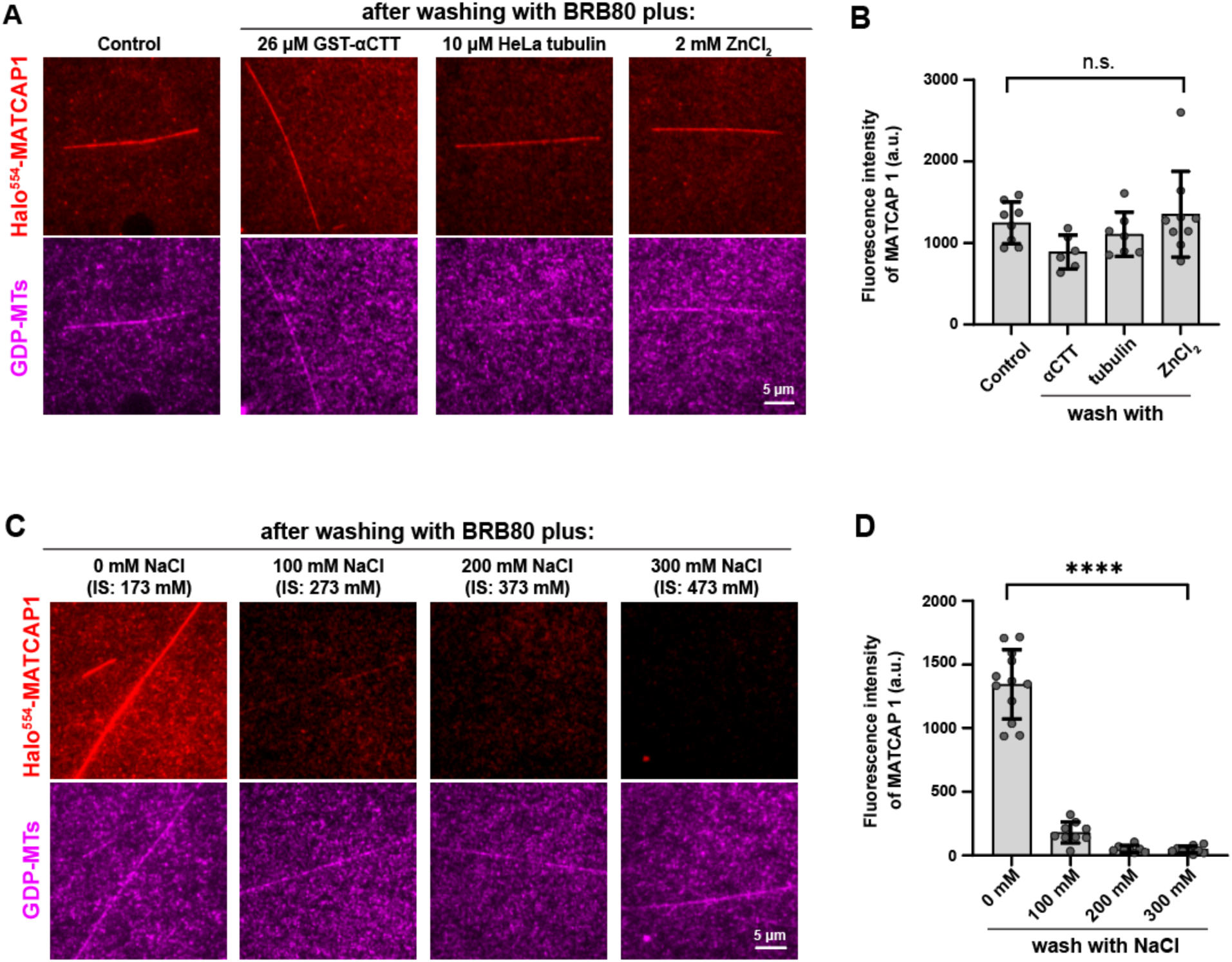
MATCAP1 detaches from microtubules in high ionic strength buffer. **(A)** Representative images of 1 nM Halo^554^-MATCAP1 (red) in cell lysates bound to glycerol-stabilized HeLa GDP-MTs (magenta) after washing with BRB80 containing 2 mM ZnCl_2_, 10 µM HeLa tubulin, or 26 µM αCTT-GST. Scale bar, 5 µm. **(B)** Quantification of the mean fluorescence intensity of Halo^554^-MATCAP1 under the indicated conditions shown in (A). Data are presented as mean ± S.D., with n = 6–9 microtubules. n.s., not significant (one-way ANOVA). **(C)** Representative images of 1 nM Halo^554^-MATCAP1 (red) in cell lysates bound to glycerol-stabilized HeLa GDP-MTs (magenta) after washing with BRB80 containing increasing concentrations of NaCl. Scale bar, 5 µm. **(D)** Quantification of the mean fluorescence intensity of Halo^554^-MATCAP1 under the indicated conditions shown in (C). Data are presented as mean ± S.D., with n = 8–12 microtubules. ****p<0.0001 (one-way ANOVA).

**Figure S5:**
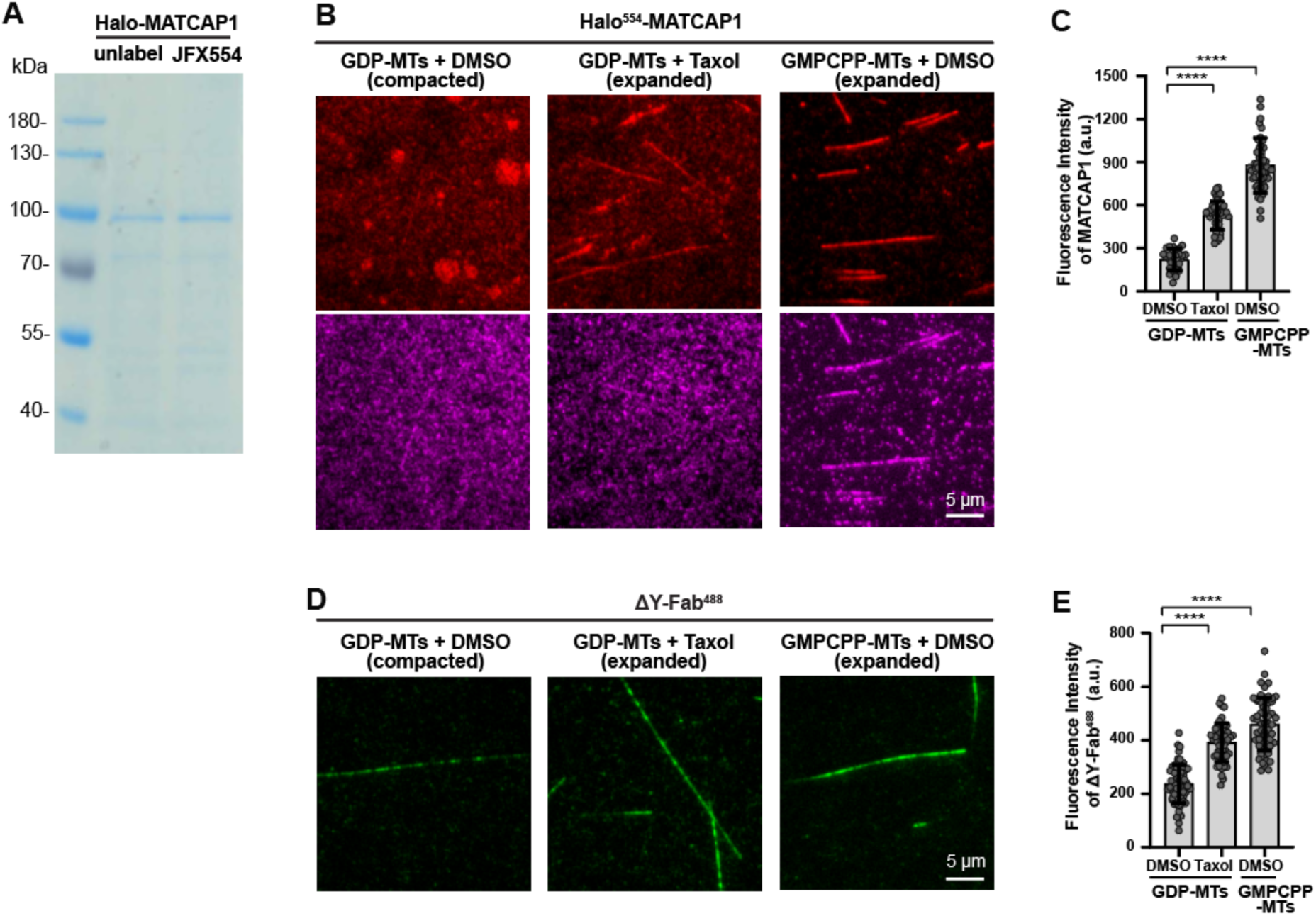
The microtubule binding and detyrosination activity of purified MATCAP1 are affected by the conformational state of the microtubule. **(A)** Coomassie-stained gel of TwinStrep-Halo-MATCAP1 protein purified from COS-7 cells. **(B,C)** Microtubule binding of purified MATCAP1 is regulated by the conformational state of tubulin in the microtubule lattice. **(B)** Representative images of 1.5 nM purified TwinStrep-Halo^554^-MATCAP1 protein binding to glycerol-stabilized GDP-MTs with DMSO, GDP-MTs with Taxol, or GMPCPP-MTs with DMSO. Scale bar, 5 µm. **(C)** Quantification of the mean fluorescence intensity of TwinStrep-Halo^554^-MATCAP1 along the microtubules in (B). Each point represents the mean fluorescence intensity of MATCAP1 along an individual microtubule. Data are presented as mean ± S.D., with n = 30–56 microtubules from two or three independent experiments. ****p<0.0001 (two-tailed, *t*-test). **(D,E)** Detyrosination activity of purified MATCAP1 is regulated by the conformational state of tubulin in the microtubule lattice. **(D)** Representative images of ΔY-Fab^488^ labeling of glycerol-stabilized GDP-MTs with DMSO, GDP-MTs with Taxol, or GMPCPP-MTs with DMSO after incubation with 1.5 nM Halo-purified MATCAP1 protein. MATCAP1 was removed by a high-salt wash step before addition of the ΔY-Fab^488^ probe. Scale bar, 5 µm. **(E)** Quantification of the mean fluorescence intensity of the ΔY-Fab^48^ probe labeling along the microtubules in (D). Each point represents the mean fluorescence intensity of ΔY-Fab^488^ along an individual microtubule. Data are presented as mean ± S.D., with n= 52–65 microtubules from two or three independent experiments. ****p<0.0001 (two-tailed, *t*-test).

## References

Aillaud, C., Bosc, C., Peris, L., Bosson, A., Heemeryck, P., Van Dijk, J., Le Friec, J., Boulan, B., Vossier, F., Sanman, L. E., Syed, S., Amara, N., Coute, Y., Lafanechere, L., Denarier, E., Delphin, C., Pelletier, L., Humbert, S., Bogyo, M., Andrieux, A., Rogowski, K. and Moutin, M. J. (2017). Vasohibins/SVBP are tubulin carboxypeptidases (TCPs) that regulate neuron differentiation. Science, 358, 1448–1453.

Aillaud, C., Bosc, C., Saoudi, Y., Denarier, E., Peris, L., Sago, L., Taulet, N., Cieren, A., Tort, O., Magiera, M. M., Janke, C., Redeker, V., Andrieux, A. and Moutin, M. J. (2016). Evidence for new C-terminally truncated variants of alpha- and beta-tubulins. Mol Biol Cell, 27, 640–653.

Alushin, G. M., Lander, G. C., Kellogg, E. H., Zhang, R., Baker, D. and Nogales, E. (2014). High-resolution microtubule structures reveal the structural transitions in alphabeta-tubulin upon GTP hydrolysis. Cell, 157, 1117–1129.

Bak, J., Brummelkamp, T. R. and Perrakis, A. (2024). Decoding microtubule detyrosination: enzyme families, structures, and functional implications. FEBS Lett, 598, 1453–1464.

Banerjee, A., Bovenzi, F. A. and Bane, S. L. (2010). High-resolution separation of tubulin monomers on polyacrylamide minigels. Anal Biochem, 402, 194–196.

Barisic, M., Silva e Sousa, R., Tripathy, S. K., Magiera, M. M., Zaytsev, A. V., Pereira, A. L., Janke, C., Grishchuk, E. L. and Maiato, H. (2015). Mitosis. Microtubule detyrosination guides chromosomes during mitosis. Science, 348, 799–803.

Berezniuk, I., Vu, H. T., Lyons, P. J., Sironi, J. J., Xiao, H., Burd, B., Setou, M., Angeletti, R. H., Ikegami, K. and Fricker, L. D. (2012). Cytosolic carboxypeptidase 1 is involved in processing alpha- and beta-tubulin. J Biol Chem, 287, 6503–6517.

Cai, D., McEwen, D. P., Martens, J. R., Meyhofer, E. and Verhey, K. J. (2009). Single molecule imaging reveals differences in microtubule track selection between Kinesin motors. PLoS Biol, 7, e1000216.

Castle, B. T., McKibben, K. M., Rhoades, E. and Odde, D. J. (2020). Tau Avoids the GTP Cap at Growing Microtubule Plus-Ends. iScience, 23, 101782.

Chen, C. Y., Caporizzo, M. A., Bedi, K., Vite, A., Bogush, A. I., Robison, P., Heffler, J. G., Salomon, A. K., Kelly, N. A., Babu, A., Morley, M. P., Margulies, K. B. and Prosser, B. L. (2018). Suppression of detyrosinated microtubules improves cardiomyocyte function in human heart failure. Nat Med, 24, 1225–1233.

Chew, Y. M. and Cross, R. A. (2025). Structural switching of tubulin in the microtubule lattice. Biochem Soc Trans, 53.

Chhatre, A., Stepanek, L., Nievergelt, A. P., Alvarez Viar, G., Diez, S. and Pigino, G. (2025). Tubulin tyrosination/detyrosination regulate the affinity and sorting of intraflagellar transport trains on axonemal microtubule doublets. Nat Commun, 16, 1055.

Cleary, J. M. and Hancock, W. O. (2021). Molecular mechanisms underlying microtubule growth dynamics. Curr Biol, 31, R560–R573.

Dunn, S., Morrison, E. E., Liverpool, T. B., Molina-Paris, C., Cross, R. A., Alonso, M. C. and Peckham, M. (2008). Differential trafficking of Kif5c on tyrosinated and detyrosinated microtubules in live cells. J Cell Sci, 121, 1085–1095.

Estevez-Gallego, J., Alvarez-Bernad, B., Pera, B., Wullschleger, C., Raes, O., Menche, D., Martinez, J. C., Lucena-Agell, D., Prota, A. E., Bonato, F., Bargsten, K., Cornelus, J., Gimenez-Abian, J. F., Northcote, P., Steinmetz, M. O., Kamimura, S., Altmann, K. H., Paterson, I., Gago, F., Van der Eycken, J., Diaz, J. F. and Oliva, M. A. (2023). Chemical modulation of microtubule structure through the laulimalide/peloruside site. Structure, 31, 88–99 e85.

Ferreira, L. T., Orr, B., Rajendraprasad, G., Pereira, A. J., Lemos, C., Lima, J. T., Guasch Boldu, C., Ferreira, J. G., Barisic, M. and Maiato, H. (2020). alpha-Tubulin detyrosination impairs mitotic error correction by suppressing MCAK centromeric activity. J Cell Biol, 219.

Gudimchuk, N. B. and McIntosh, J. R. (2021). Regulation of microtubule dynamics, mechanics and function through the growing tip. Nat Rev Mol Cell Biol, 22, 777–795.

Hotta, T., Plemmons, A., Gebbie, M., Ziehm, T. A., Blasius, T. L., Johnson, C., Verhey, K. J., Pearring, J. N. and Ohi, R. (2023). Mechanistic Analysis of CCP1 in Generating DeltaC2 alpha-Tubulin in Mammalian Cells and Photoreceptor Neurons. Biomolecules, 13.

Huzil, J. T., Chik, J. K., Slysz, G. W., Freedman, H., Tuszynski, J., Taylor, R. E., Sackett, D. L. and Schriemer, D. C. (2008). A unique mode of microtubule stabilization induced by peloruside A. J Mol Biol, 378, 1016–1030.

Janke, C. and Magiera, M. M. (2020). The tubulin code and its role in controlling microtubule properties and functions. Nat Rev Mol Cell Biol, 21, 307–326.

Khandelia, P., Yap, K. and Makeyev, E. V. (2011). Streamlined platform for short hairpin RNA interference and transgenesis in cultured mammalian cells. Proc Natl Acad Sci U S A, 108, 12799–12804.

Konietzny, A., Han, Y., Popp, Y., van Bommel, B., Sharma, A., Delagrange, P., Arbez, N., Moutin, M. J., Peris, L. and Mikhaylova, M. (2024). Efficient axonal transport of endolysosomes relies on the balanced ratio of microtubule tyrosination and detyrosination. J Cell Sci, 137.

Konishi, Y. and Setou, M. (2009). Tubulin tyrosination navigates the kinesin-1 motor domain to axons. Nat Neurosci, 12, 559–567.

LaFrance, B. J., Roostalu, J., Henkin, G., Greber, B. J., Zhang, R., Normanno, D., McCollum, C. O., Surrey, T. and Nogales, E. (2022). Structural transitions in the GTP cap visualized by cryo-electron microscopy of catalytically inactive microtubules. Proc Natl Acad Sci U S A, 119.

Landskron, L., Bak, J., Adamopoulos, A., Kaplani, K., Moraiti, M., van den Hengel, L. G., Song, J. Y., Bleijerveld, O. B., Nieuwenhuis, J., Heidebrecht, T., Henneman, L., Moutin, M. J., Barisic, M., Taraviras, S., Perrakis, A. and Brummelkamp, T. R. (2022). Posttranslational modification of microtubules by the MATCAP detyrosinase. Science, 376, eabn6020.

Lavrsen, K., Rajendraprasad, G., Leda, M., Eibes, S., Vitiello, E., Katopodis, V., Goryachev, A. B. and Barisic, M. (2023). Microtubule detyrosination drives symmetry breaking to polarize cells for directed cell migration. Proc Natl Acad Sci U S A, 120, e2300322120.

Li, F., Li, Y., Ye, X., Gao, H., Shi, Z., Luo, X., Rice, L. M. and Yu, H. (2020). Cryo-EM structure of VASH1-SVBP bound to microtubules. Elife, 9.

Liao, S., Rajendraprasad, G., Wang, N., Eibes, S., Gao, J., Yu, H., Wu, G., Tu, X., Huang, H., Barisic, M. and Xu, C. (2019). Molecular basis of vasohibins-mediated detyrosination and its impact on spindle function and mitosis. Cell Res, 29, 533–547.

Liu, H. and Shima, T. (2023). Preference of CAMSAP3 for expanded microtubule lattice contributes to stabilization of the minus end. Life Sci Alliance, 6.

Manka, S. W. and Moores, C. A. (2018). Microtubule structure by cryo-EM: snapshots of dynamic instability. Essays Biochem, 62, 737–751.

McKenney, R. J., Huynh, W., Vale, R. D. and Sirajuddin, M. (2016). Tyrosination of alpha-tubulin controls the initiation of processive dynein-dynactin motility. EMBO J, 35, 1175–1185.

Moutin, M. J., Bosc, C., Peris, L. and Andrieux, A. (2021). Tubulin post-translational modifications control neuronal development and functions. Dev Neurobiol, 81, 253–272.

Nicot, S., Gillard, G., Impheng, H., Joachimiak, E., Urbach, S., Mochizuki, K., Wloga, D., Juge, F. and Rogowski, K. (2023). A family of carboxypeptidases catalyzing alpha- and beta-tubulin tail processing and deglutamylation. Sci Adv, 9, eadi7838.

Nieuwenhuis, J., Adamopoulos, A., Bleijerveld, O. B., Mazouzi, A., Stickel, E., Celie, P., Altelaar, M., Knipscheer, P., Perrakis, A., Blomen, V. A. and Brummelkamp, T. R. (2017). Vasohibins encode tubulin detyrosinating activity. Science, 358, 1453–1456.

Nieuwenhuis, J. and Brummelkamp, T. R. (2019). The Tubulin Detyrosination Cycle: Function and Enzymes. Trends Cell Biol, 29, 80–92.

Nirschl, J. J., Magiera, M. M., Lazarus, J. E., Janke, C. and Holzbaur, E. L. (2016). alpha-Tubulin Tyrosination and CLIP-170 Phosphorylation Regulate the Initiation of Dynein-Driven Transport in Neurons. Cell Rep, 14, 2637–2652.

Paquette, A. L., Cruz Tetlalmatzi, S., Haineault, J. A. G., Hendricks, A. G., Brouhard, G. J. and Sébastien, M. (2025). Competition for microtubule lattice spacing between a microtubule expander and compactor. bioRxiv.

Peet, D. R., Burroughs, N. J. and Cross, R. A. (2018). Kinesin expands and stabilizes the GDP-microtubule lattice. Nat Nanotechnol, 13, 386–391.

Peris, L., Wagenbach, M., Lafanechere, L., Brocard, J., Moore, A. T., Kozielski, F., Job, D., Wordeman, L. and Andrieux, A. (2009). Motor-dependent microtubule disassembly driven by tubulin tyrosination. J Cell Biol, 185, 1159–1166.

Prota, A. E., Lucena-Agell, D., Ma, Y., Estevez-Gallego, J., Li, S., Bargsten, K., Josa-Prado, F., Altmann, K. H., Gaillard, N., Kamimura, S., Muhlethaler, T., Gago, F., Oliva, M. A., Steinmetz, M. O., Fang, W. S. and Diaz, J. F. (2023). Structural insight into the stabilization of microtubules by taxanes. Elife, 12.

Ramirez-Rios, S., Choi, S. R., Sanyal, C., Blum, T. B., Bosc, C., Krichen, F., Denarier, E., Soleilhac, J. M., Blot, B., Janke, C., Stoppin-Mellet, V., Magiera, M. M., Arnal, I., Steinmetz, M. O. and Moutin, M. J. (2023). VASH1-SVBP and VASH2-SVBP generate different detyrosination profiles on microtubules. J Cell Biol, 222.

Robison, P., Caporizzo, M. A., Ahmadzadeh, H., Bogush, A. I., Chen, C. Y., Margulies, K. B., Shenoy, V. B. and Prosser, B. L. (2016). Detyrosinated microtubules buckle and bear load in contracting cardiomyocytes. Science, 352, aaf0659.

Rogowski, K., van Dijk, J., Magiera, M. M., Bosc, C., Deloulme, J. C., Bosson, A., Peris, L., Gold, N. D., Lacroix, B., Bosch Grau, M., Bec, N., Larroque, C., Desagher, S., Holzer, M., Andrieux, A., Moutin, M. J. and Janke, C. (2010). A family of protein-deglutamylating enzymes associated with neurodegeneration. Cell, 143, 564–578.

Roll-Mecak, A. (2020). The Tubulin Code in Microtubule Dynamics and Information Encoding. Dev Cell, 54, 7–20.

Salomon, A. K., Phyo, S. A., Okami, N., Heffler, J., Robison, P., Bogush, A. I. and Prosser, B. L. (2022). Desmin intermediate filaments and tubulin detyrosination stabilize growing microtubules in the cardiomyocyte. Basic Res Cardiol, 117, 53.

Sanyal, C., Pietsch, N., Ramirez Rios, S., Peris, L., Carrier, L. and Moutin, M. J. (2023). The detyrosination/re-tyrosination cycle of tubulin and its role and dysfunction in neurons and cardiomyocytes. Semin Cell Dev Biol, 137, 46–62.

Schmidt, T. G., Batz, L., Bonet, L., Carl, U., Holzapfel, G., Kiem, K., Matulewicz, K., Niermeier, D., Schuchardt, I. and Stanar, K. (2013). Development of the Twin-Strep-tag(R) and its application for purification of recombinant proteins from cell culture supernatants. Protein Expr Purif, 92, 54–61.

Shen, Y. and Ori-McKenney, K. M. (2024). Microtubule-associated protein MAP7 promotes tubulin posttranslational modifications and cargo transport to enable osmotic adaptation. Dev Cell, 59, 1553–1570 e1557.

Shima, T., Morikawa, M., Kaneshiro, J., Kambara, T., Kamimura, S., Yagi, T., Iwamoto, H., Uemura, S., Shigematsu, H., Shirouzu, M., Ichimura, T., Watanabe, T. M., Nitta, R., Okada, Y. and Hirokawa, N. (2018). Kinesin-binding-triggered conformation switching of microtubules contributes to polarized transport. J Cell Biol, 217, 4164–4183.

Siahaan, V., Tan, R., Humhalova, T., Libusova, L., Lacey, S. E., Tan, T., Dacy, M., Ori-McKenney, K. M., McKenney, R. J., Braun, M. and Lansky, Z. (2022). Microtubule lattice spacing governs cohesive envelope formation of tau family proteins. Nat Chem Biol, 18, 1224–1235.

Souphron, J., Bodakuntla, S., Jijumon, A. S., Lakisic, G., Gautreau, A. M., Janke, C. and Magiera, M. M. (2019). Purification of tubulin with controlled post-translational modifications by polymerization-depolymerization cycles. Nat Protoc, 14, 1634–1660.

Thomas, E. C., Yue, Y., Pimm, M. L., Hotta, T., Ohi, R. and Verhey, K. J. (2025). Purification, fluorescent labelling, and detyrosination of mammalian cell tubulin for biochemical assays. Cytoskeleton (Hoboken).

Tort, O., Tanco, S., Rocha, C., Bieche, I., Seixas, C., Bosc, C., Andrieux, A., Moutin, M. J., Aviles, F. X., Lorenzo, J. and Janke, C. (2014). The cytosolic carboxypeptidases CCP2 and CCP3 catalyze posttranslational removal of acidic amino acids. Mol Biol Cell, 25, 3017–3027.

Verhey, K. J. and Gaertig, J. (2007). The tubulin code. Cell Cycle, 6, 2152–2160.

Yue, Y., Hotta, T., Higaki, T., Verhey, K. J. and Ohi, R. (2023). Microtubule detyrosination by VASH1/SVBP is regulated by the conformational state of tubulin in the lattice. Curr Biol, 33, 4111–4123 e4117.

Zhang, R., LaFrance, B. and Nogales, E. (2018). Separating the effects of nucleotide and EB binding on microtubule structure. Proc Natl Acad Sci U S A, 115, E6191–E6200.

